# Genome-wide transcription factor binding in leaves from C_3_ and C_4_ grasses

**DOI:** 10.1101/165787

**Authors:** Steven J. Burgess, Ivan Reyna-Llorens, Sean R. Stevenson, Pallavi Singh, Katja Jaeger, Julian M. Hibberd

## Abstract

The majority of plants use C_3_ photosynthesis, but over sixty independent lineages of angiosperms have evolved the C_4_ pathway. In most C_4_ species, photosynthesis gene expression is compartmented between mesophyll and bundle sheath cells. We performed DNaseI-SEQ to identify genome-wide profiles of transcription factor binding in leaves of the C_4_ grasses *Zea mays*, *Sorghum bicolor* and *Setaria italica* as well as C_3_ *Brachypodium distachyon*. In C_4_ species, while bundle sheath strands and whole leaves shared similarity in the broad regions of DNA accessible to transcription factors, the short sequences bound varied. Transcription factor binding was prevalent in gene bodies as well as promoters, and many of these sites could represent duons that impact gene regulation in addition to amino acid sequence. Although globally there was little correlation between any individual DNaseI footprint and cell-specific gene expression, within individual species transcription factor binding to the same motifs in multiple genes provided evidence for shared mechanisms governing C_4_ photosynthesis gene expression. Furthermore, interspecific comparisons identified a small number of highly conserved transcription factor binding sites associated with leaves from species that diverged around 60 million years ago. These data therefore provide insight into the architecture associated with C_4_ photosynthesis gene expression in particular and characteristics of transcription factor binding in cereal crops in general.

**One sentence summary:** Genome-wide patterns of transcription factor binding *in vivo* defined by DNaseI for leaves of C_3_ and C_4_ grasses

## Introduction

Most photosynthetic organisms, including crops of global importance such as wheat, rice and potato use the C_3_ photosynthesis pathway in which Ribulose-Bisphosphate Carboxylase Oxygenase (RuBisCO) catalyses the primary fixation of CO_2_. However, carboxylation by RuBisCO is competitively inhibited by oxygen binding the active site (Bowes et al., 1971). This oxygenation reaction generates toxic waste-products that are recycled by an energy-demanding series of metabolic reactions known as photorespiration (Bauwe et al., 2010; Tolbert, 1971). The ratio of oxygenation to carboxylation increases with temperature (Jordan and Ogren, 1984; Sharwood et al., 2016) and so losses from photorespiration are particularly high in the tropics.

Multiple plant lineages have evolved mechanisms that suppress oxygenation by concentrating CO_2_ around RuBisCO. One such strategy is known as C_4_ photosynthesis. Species that use the C_4_ pathway include maize, sorghum and sugarcane, and they represent the most productive crops on the planet (Sage and Zhu, 2011). In C_4_ leaves, additional expenditure of ATP, alterations to leaf anatomy and cellular ultrastructure, as well as spatial separation of photosynthesis between compartments (Hatch, 1987) allows CO_2_ concentration to be increased around tenfold compared with that in the atmosphere (Furbank, 2011). Despite the complexity of C_4_ photosynthesis, it is found in over 60 independent plant lineages (Sage et al., 2011). In most C_4_ plants the initial RuBisCO-independent fixation of CO_2_ and the subsequent RuBisCO-dependent reactions take place in distinct cell-types known as mesophyll and bundle sheath cells. Although the spatial patterning of gene expression that generates these metabolic specialisations is fundamental to C_4_ photosynthesis very few examples of *cis*-elements or *trans*-factors that restrict gene expression to mesophyll or bundle sheath cells of C_4_ plants have been identified (Brown et al., 2011; Gowik et al., 2004; Williams et al., 2016; Reyna-Llorens et al., 2018). Moreover, in grasses more generally the DNA-binding properties of relatively few transcription factors have been validated (Bolduc and Hake, 2009; Yu et al., 2015; Eveland et al., 2014; Pautler et al., 2015). In summary, in both C_3_ and C_4_ species, work has focussed on analysis of mechanisms controlling the expression of individual genes, and so our understanding of the overall landscape associated with photosynthesis gene expression is poor.

In yeast and animal systems, the high sensitivity of open chromatin to DNaseI (Zentner and Henikoff, 2014) has allowed comprehensive, genome-wide characterization of transcription factor binding sites at single nucleotide resolution (Hesselberth et al., 2009; Neph et al., 2012; Thurman et al., 2012). In plants, DNaseI-SEQ and more recently Assay for Transposase-Accessible Chromatin (ATAC-SEQ) have been employed in C_3_ species and provided insight into the patterns of transcription factor binding associated with development (Zhang et al., 2012a; Pajoro et al., 2014; Zhang et al., 2012b, 2016), heat stress (Sullivan et al., 2014) and root cell differentiation (Maher et al., 2017). By carrying out DNaseI-SEQ on grass leaves that use either C_3_ or C_4_ photosynthesis, we aimed to provide insight into the transcription factor binding repertoire associated with each form of photosynthesis. Our data indicate more transcription factor binding sites are found in gene bodies than promoters, and up to 25% of the footprints represent ‘duons’ – sequences located in exons that have an impact on both gene regulation as well as the amino acid sequence of the protein they encode. It is also clear that specific cell types from leaf tissue make use of a markedly distinct *cis*-regulatory code and that despite significant turnover in the cistrome of grasses, a small number of transcription factor motifs are conserved across 60 million years of evolution. Comparison of sites bound by transcription factors in both C_3_ and C_4_ leaves demonstrates that the repeated evolution of C_4_ photosynthesis is built on both the *de novo* gain of *cis*-elements and the exaptation of highly conserved regulatory elements found in the ancestral C_3_ system.

## Results

### A *cis*-regulatory atlas for grasses

To provide insight into the regulatory architecture associated with C_3_ and C_4_ photosynthesis in cereal crops, four grass were selected. *Brachypodium distachyon* uses the ancestral C_3_ pathway (Figure 1A). *Sorghum bicolor, Zea mays* and *Setaria italica* all use C_4_ photosynthesis, they were chosen as phylogenetic reconstructions indicate that *S. italica* represents an independent evolutionary origin of the C_4_ pathway (Figure 1A) and comparison of these species can provide insight into parallel and convergent evolution of C_4_ gene expression. Nuclei from a minimum of duplicate samples of *S. italica* (C_4_), *S. bicolor* (C_4_), *Z. mays* (C_4_) and *B. distachyon* (C_3_) leaves were treated with DNaseI (Supplemental Figure 1) and subjected to deep sequencing. A total of 806,663,951 reads could be uniquely mapped to the respective genome sequences of these species (Supplemental Table 1). From all four genomes, 159,396 DNaseI-hypersensitive sites (DHS) of between 150-15,060 base pairs representing broad regulatory regions accessible to transcription factor binding were identified (Figure 1B). Between 20,817 and 27,746 genes were annotated as containing at least one DHS (Supplemental Table 2). For subsequent analysis, only DHS that were consistent between replicates as determined by the Irreproducible Discovery Rate framework (Li and Dewey, 2011) were used.

**Figure 1:**
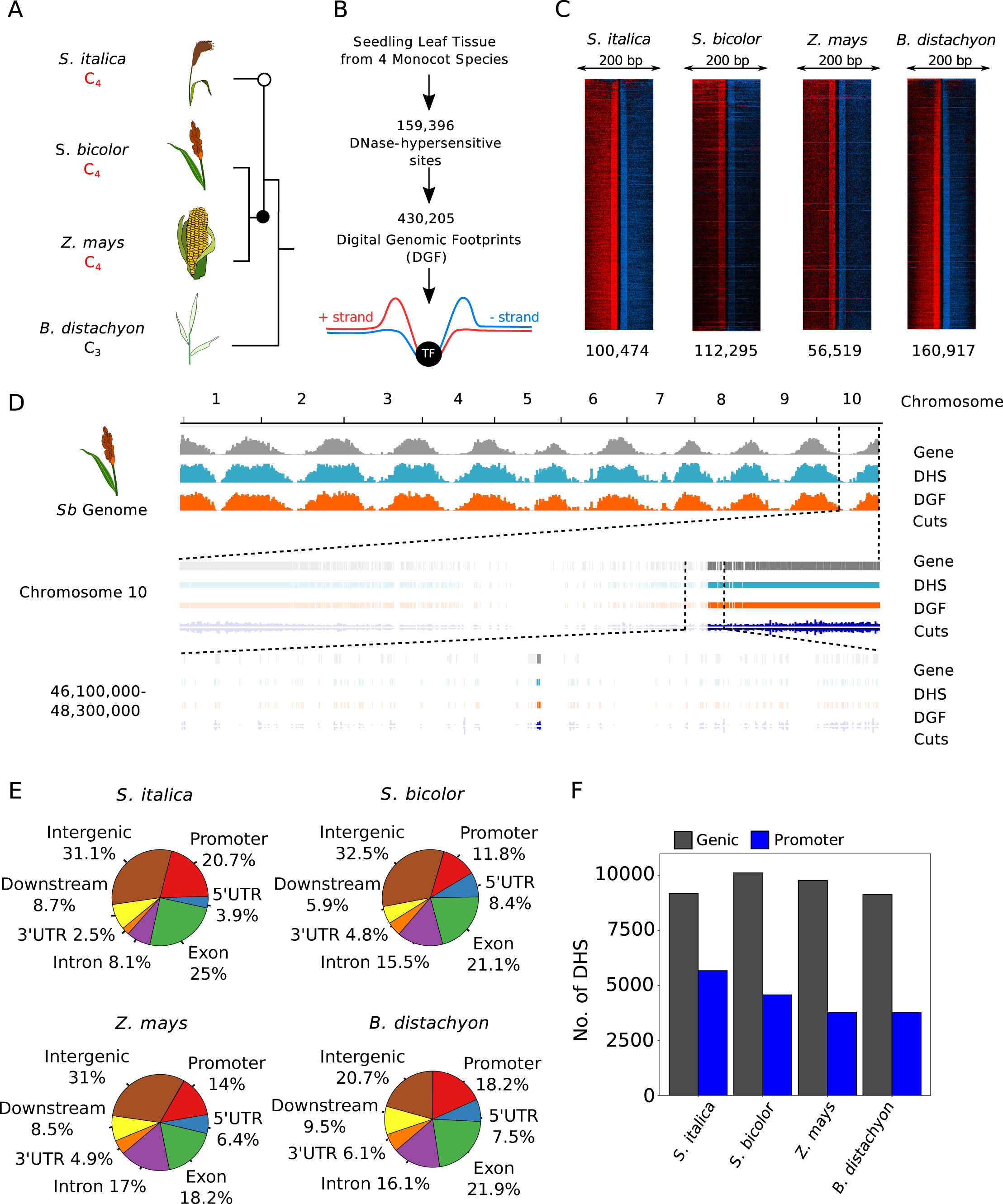
Transcription factor binding atlas for whole leaf samples of four grasses. Schematic of phylogenetic relationship between species analysed. The two independent origins of C_4_ photosynthesis are highlighted with black and white circles (figure not drawn to scale). (B) Summary of sampling and the total number of DNaseI-hypersensitive sites (DHS) and Digital Genomic Footprints (DGF) identified across all four species. (C) TreeView diagrams illustrating cut density around individual digital genomic footprint (DGF). Each row represents an individual DGF, cuts are coloured according to whether they align to the positive (red) or negative (blue) strand and indicate increased cutting in a 50 bp window on either side of the DGF. The total number of DGF per sample is shown at the bottom. (D) Representation of DNaseI-SEQ data from *S. bicolor*, depicting gene (grey), DHS (light blue), DGF (orange) and DNaseI cut density (dark blue) at three scales: genome wide, with chromosome number and position indicated (top), chromosomal (second level) and kilobase genomic region (third level). Between each level the expanded area is denoted by dashed lines. (E) Pie-chart representing the distribution of DHS among genomic features. Promoters are defined as sequence up to 2000 base pairs (bp) upstream of the transcriptional start site, downstream represent regions 1000 downstream the transcription termination site while intergenic represent > 1000 bp downstream the transcription termination site until the next promoter region. (F) Bar chart representing number of DHS in genic and promoter regions.

DNaseI footprinting is a well-established technique for detecting DNA-protein interactions at base pair resolution and as such has been used to generate Digital Genomic Footprints (DGF) to predict transcription factor binding sites. DGF are obtained by pooling all replicates to maximise the number of reads that map within each DHS, and then modelling differential accumulation of reads mapping to positive or negative strands around transcription factor binding sites within the DHS (Piper et al., 2013). However, the DNaseI enzyme possesses some sequence bias that can affect prediction of transcription factor binding sites (He et al., 2014; Yardimci et al., 2014). After performing DNaseI-SEQ on “naked DNA” that is devoid of nucleosomes from each species, we identified hundreds of DGF that likely represent false positives (Supplemental Figure 2A). For all species, analysis of the DGF derived from naked DNA showed that treatment with DNaseI led to similar sequences being preferentially digested (Supplemental Figure 2B). However, because false positive DGF predicted from this approach will be influenced by the number of reads that map to each genome, and in the case of maize fewer reads mapped in total, the number of false positive DGF varied between species (Supplemental Figure 2A). To overcome this issue, we implemented a more conservative pipeline that rather than defining false positives at specific locations within the genome, calculates DNaseI cutting bias for all hexamers across each genome. By employing a mixture model framework, these data are then used to generate a background signal to estimate footprint likelihood scores for each putative DGF (Yardimici et al., 2014; Supplemental Figure 2B). This approach removed between 15% and 30% of DGF from each sample (Supplemental Figure 2C) and left a total of 430,205 DGF corresponding to individual transcription factor binding sites between 11 and 25 base pairs being identified (Figure 1B&C; Supplemental Table 3). At least one transcription factor footprint was identified in >75% of the broader regions defined by DHS (Supplemental Table 2).

We attempted to saturate the number of predicted DGF by sequencing each species at high depth (Supplemental Table 1). *In silico* sub-sampling of these data indicated that for *S. bicolor, S. italica* and *B. distachyon*, the total number of DGF was close to saturation, but for maize despite obtaining 251,955,063 reads from whole leaves this was not the case (Supplementary Figure 3). Consequently, fewer DGF were predicted in maize compared with the other species (Figure 1C). Since maize has a similar gene number to the other species analysed, it is possible that the reduced ability to map reads to unique loci was associated with the high amount of repetitive DNA in the maize genome. Another contributing factor to the poor mapping rate in maize may be the low complexity found in one of the libraries as reflected by the PCR bottleneck coefficient (Supplemental Table 1). According to the Encyclopaedia of DNA elements (ENCODE), large number of reads from low complexity libraries decreases the chances of identifying the majority of transcription factor binding sites. However, despite these differences in coverage and in certain quality metrics, for all four species DHS and DGF were primarily located in gene-rich regions and depleted around centromeres (Figure 1D). Individual transcription factor binding sequences were resolved in all chromosomes from each species (Figure 1D). On a genome-wide basis, the distribution of DHS was similar between species, with the highest proportion of such sites located in promoter, coding sequence and intergenic regions (Figure 1E). Notably, in all four grasses, genic sequences contained more DHS than promoters (Figure 1F).

To provide additional evidence confirming that DGF identified in our analysis derive from protein-DNA interactions, they were compared with previously identified motifs from maize. Maize is the most appropriate choice for this analysis as there are more data on transcription factor binding sites than in *S. bicolor* and *S. italica*. Moreover, support from previous work goes some way to supporting the smaller number of DGF that we identified in this species. Therefore, the literature was assessed for validated transcription factor binding sites in maize. These have previously been associated with flowering (Kozaki et al., 2004; Vollbrecht et al., 2005; Eveland et al., 2014), meristem development (Bolduc et al., 2012), Gibberellin catabolism (Bolduc and Hake, 2009), sugar signalling (Niu et al., 2002) and leaf development of maize (Yu et al., 2015), but in all cases, DGF matching these motifs were found in our dataset (Figure 2A, FDR < 0.001). In addition, a larger ChIP-SEQ dataset of 117 transcription factors from maize leaves obtained from the pre-release maize cistrome (http://www.epigenome.cuhk.edu.hk/C3C4.html, Supplemental Figure 4, Supplemental Table 4) was compared with our data. Differences between specific binding sites are likely because in all cases growth conditions will have varied from ours, and in some cases different tissues were sampled. Despite this, 66% and 29% of the ChIP-SEQ peaks overlapped with our DHS and DGF respectively. Although only 29% of DGF overlapped with motifs defined by ChIP-SEQ, permutation tests performed using the “regioneR” package (Gel et al., 2015) indicated a statistically greater overlap than would be expected by chance (pvalue = 0.0099, 100 permutations). Moreover, when both features were systematically shifted from their original position, the local z-score, which represents the strength of the association at any particular position showed a sharper decrease for DGF than DHS suggesting the association between ChIP-SEQ peaks and DGFs is more strongly linked to the exact position of the DGF (Supplemental Figure 4B). In summary, despite detecting fewer DGF in maize than in the other species, the DGF we found are supported by publicly available ChIP-SEQ, EMSA and Selex datasets.

**Figure 2:**
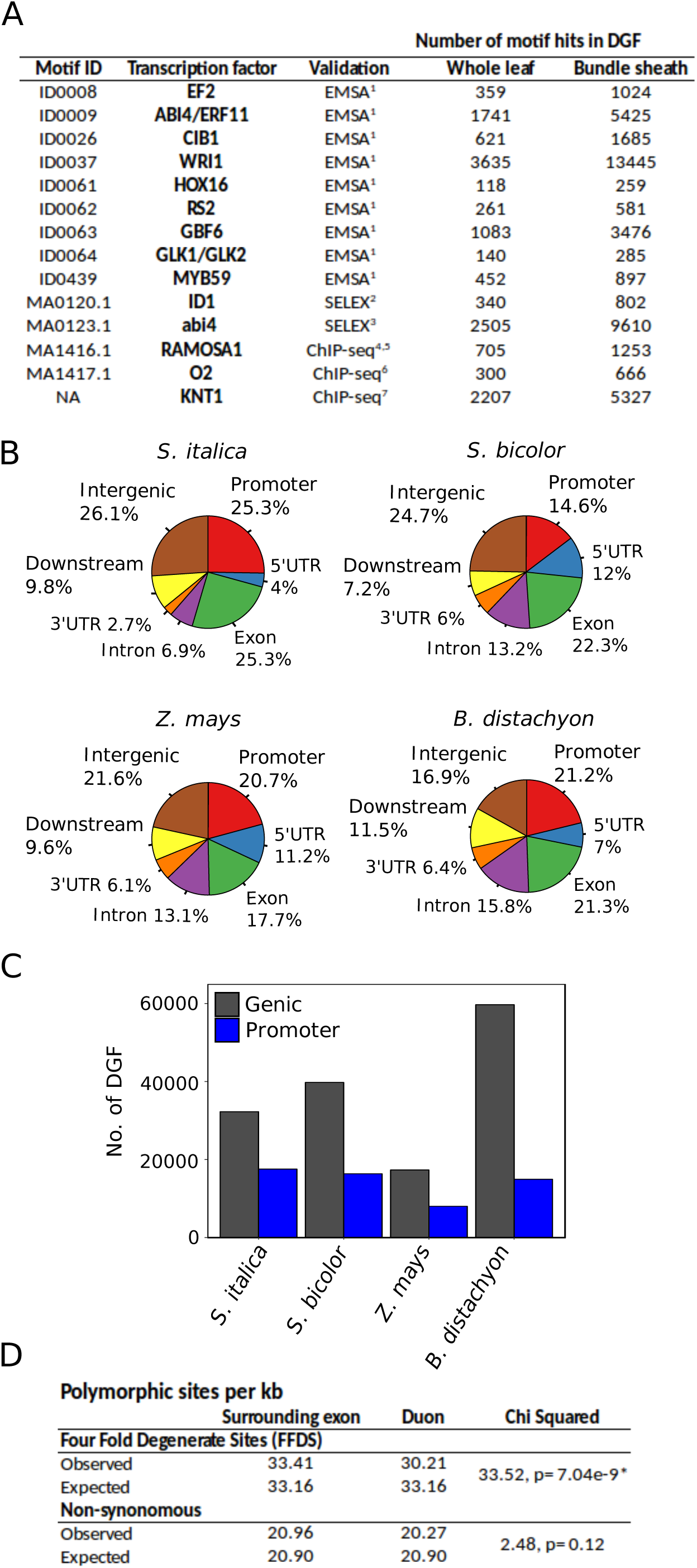
Digital genomic footprints in whole leaves of four grasses. (A) DNA motifs from previous studies in maize (^1^Yu et al., 2015; ^2^Kozaki et al., 2004; ^3^Niu et al., 2002; ^4^Vollbrecht et al., 2005; ^5^Eveland et al., 2014; ^6^Li et al., 2015; ^7^Bolduc et al., 2012) were detected in whole leaves and bundle sheath strands from maize. (B) Pie-chart representing the distribution of DGF among genomic features. Promoters are defined as sequence up to 2000 base pairs (bp) upstream of the transcriptional start site, downstream represent regions 1000 downstream the transcription termination site while intergenic represent > 1000 bp downstream the transcription termination site until the next promoter region. (C) Bar chart representing number of DGF in genic and promoter regions. (D) Polymorphic sites per kb in duons and surrounding exons at FourFold Degenerate Sites (FFDS) and non-synonymous sites. Chi-squared tests indicate reduced rates of mutation at FFDS than expected by chance.

Consistent with the distribution of DHS (Figure 1E), annotated DGF were most common in promoter, coding sequence and intergenic regions (Figure 2B) and genic sequences contained more DGF than promoters (Figure 2C). Distribution plots showed that the highest density of DGF was close to the annotated transcription start sites but indicated a slightly skewed distribution favouring genic sequence including exons (Supplemental Figure 5). A similar pattern was observed for the ChIP-SEQ signal peaks (Supplemental Figure 4C). Transcription factor binding sites located in exons have been termed duons because they could impact both on the regulation of transcription and amino acid sequence. Whilst in general synonymous mutations not affecting amino acid sequence should be under relaxed purifying selection, because of transcription factor recognition all nucleotides in duons should be under purifying selection, and thus show lower mutation rates. We therefore investigated the nucleotide substitution rate at Four-Fold Degenerate Sites (FFDS) using variation data from 1218 maize lines (Bukowski et al., 2018) and found that it was statistically significantly lower in duons than in surrounding coding sequence (Figure 2E, p= 7.04e-9). This contrasts with the density of polymorphisms in non-synonymous sites (Figure 2D). Although it has been proposed that GC bias of duons constrains FFDS (Xing and He, 2014) we found no such bias between duon and exon sequences used in this analysis (Supplemental Figure 6). Taken together, we conclude that in these cereals a significant proportion of transcription factor binding likely takes place within genes.

### A distinct *cis*-regulatory lexicon for specific cells within the leaf

The above analysis provides a genome-wide overview of the *cis*-regulatory architecture associated with leaves of grasses. However, as with other complex multicellular systems, leaves are composed of many specialised cell types. Because DGF are defined by the differential DNA cleavage between protected and unprotected regions of DNA within a DHS, a negative distribution compared with the larger DHS is produced (Figure 3A). Thus, transcription factor binding signal from a low abundance cell-type is likely to be obscured by overall signal from a tissue-level analysis (Figure 3A). Since bundle sheath strands can be separated (Covshoff et al., 2013; Leegood, 1985; Furbank et al., 1985) C_4_ species provide a simple system to study transcription factor binding in specific cells of leaves (Figure 3B). After bundle sheath isolation from *S. bicolor, S. italica* and *Z. mays*, and naked DNA correction for inherent bias in DNaseI cutting, a total of 129,137 DHS were identified (Figure 3B; Supplemental Table 5) containing 244,554 DGF (Figure 3B; Supplemental Table 5; FDR<0.01). Of these, 138,075 were statistically enriched in the bundle sheath samples compared with whole leaves (Figure 3B; Supplemental Table 5). The number of these statistically enriched DGF in bundle sheath strands of C_4_ species was large and ranged from 14,250 to 73,057 in maize and *S. italica* respectively (Supplemental Table 5). The lower number in maize is likely due to the reduced sequencing depth achieved. Genome-wide, the number of broad regulatory regions defined by DHS in the bundle sheath that overlapped with those present in whole leaves ranged from 71 to 84% in *S. italica* and *S. bicolor* respectively (Supplemental Table 6). However, only 6-20% of the narrower DGF found in the bundle sheath were also identified in whole leaves (Supplemental Table 7). Taken together, these findings indicate that specific cell types of cereal leaves share similarity in the broad regions of DNA that are accessible to transcription factors (DHS), but that the short sequences actually bound by transcription factors (DGF) vary dramatically.

**Figure 3:**
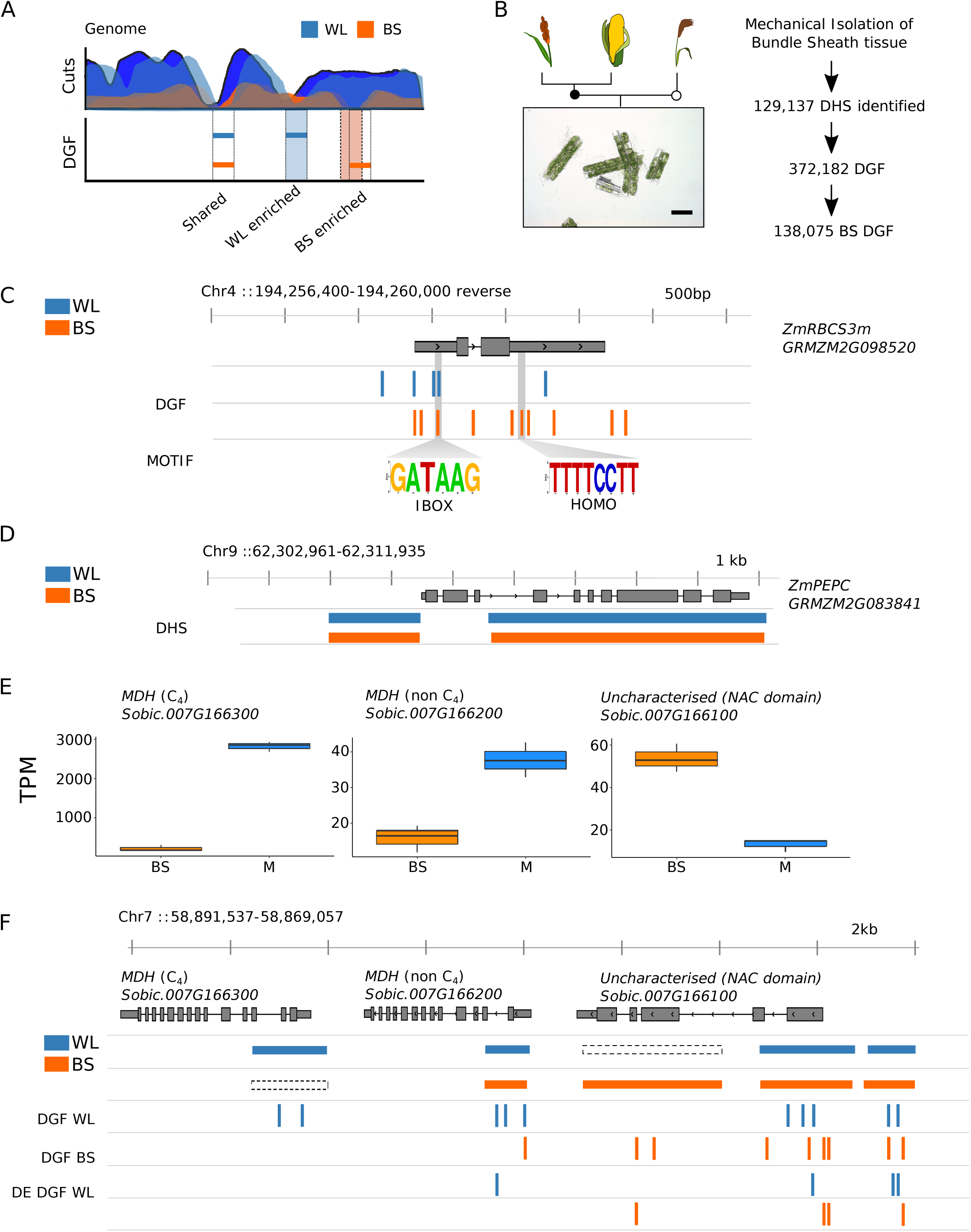
Characterisation of the DNA binding landscape in the C_4_ Bundle Sheath. (A) Schematic showing that due to their negative distribution below the background signal derived from reads mapping to the genome, footprints associated with low abundance cells such as the Bundle Sheath (BS) are unlikely to be detected from whole leaf (WL) samples. (B) Bundle sheath isolation for DNase I-SEQ experiments, with phylogeny (left) and workflow (right). (C) DGF identified in the maize *ZmRBCS3* gene coincide with I- and HOMO-boxes known to regulate gene expression. The gene model is annotated with whole leaf (blue) and BS (orange) DGF, and the I- and HOMO-boxes are indicated below. (D) DHS distribution across the maize *PEPC* gene in BS and WL samples. (E) Transcript abundance expressed as transcripts per million reads (TPM) of three co-linear genes on chromosome seven of sorghum - C_4_ *MDH* (Sobic.007G166300), non C_4_ MDH (non C_4_, Sobic.007G166200) and an uncharacterised NAC domain protein (Sobic.007G166100) in bundle sheath and mesophyll cells. Schematic of these co-linear genes from *S. bicolor*, depicting three classes of alterations to DNA accessibility and transcription factor binding to genes that are differentially expressed between whole leaf and bundle sheath cells. (F) Whole leaf (blue) and bundle sheath (orange) DHS, DGF and differentially (DE) enriched DGF, as determined by the Wellington Bootstrap algorithm, are depicted. Regions where a DHS was identified in one sample but not another are indicated by dashed boxes.

To provide evidence that DGF predicted after analysis of separated bundle sheath strands are of functional importance, they were compared with previously validated sequences. In C_4_ grasses, to our knowledge there are no such examples in *S. bicolor* or *S. italica*, but in the *RbcS* gene from maize, which is preferentially expressed in bundle sheath cells, an I-box (GATAAG) is essential for light-mediated activation (Giuliano et al., 1988) and a HOMO motif (CCTTTTTTCCTT) is important in driving bundle sheath expression (Xu et al., 2001) (Figure 3C). Despite not reaching saturation in DGF prediction in maize (Supplemental Figure 3) both elements were detected in our pipeline. Interestingly, a signal suggesting TF binding to the HOMO motif was enriched in the bundle sheath strands (Figure 3C), and whilst the I-box was detected in both bundle sheath strands and whole leaves its position was slightly different in each cell type (Figure 3C). These findings are therefore consistent with the biochemical data implicating the I-box in control of abundance and the HOMO box in control of cell-specific accumulation of *RbcS* transcripts.

The *ZmPEPC* gene (GRMZM2G083841) encodes the phospho*enol*pyruvate carboxylase responsible for producing C_4_ acids used in the C_4_ pathway and is preferentially expressed in mesophyll cells. Previous reports showed that a region of 600 nucleotides upstream of the transcription start site carrying repeated C-rich sequences was sufficient to drive expression in mesophyll cells of maize (Shaffner & Sheen, 1992; Matzuoka et al., 1994). Although no DGF were detected with these C-rich sequences, they are located within a DHS indicating that they are available for transcription factor binding (Figure 3D). Thus, despite the fact that we had not reached saturation of DGF in maize, for both *RbcS* and *PEPC* the regions of DNA accessible to transcription factor binding are consistent with previous reports, and in the case of *RbcS* DGF were detected that coincide with known *cis*-elements. To investigate the relationship between cell specific gene expression and the position of DHS and DGF, the DNaseI data were interrogated using RNA-SEQ datasets from mesophyll and bundle sheath cells of C_4_ leaves (Chang et al., 2012; John et al., 2014; Emms et al., 2016). At least three mechanisms associated with cell specific gene expression operating around individual genes were identified and can be exemplified using three co-linear genes found on chromosome seven of *S. bicolor*. First, in the *NADP-malate dehydrogenase* (*MDH*) gene, which is highly expressed in mesophyll cells and encodes a protein of the core C_4_ cycle (Figure 3E) a broad DHS site and two DGF were present in whole leaves, but not in bundle sheath strands (Figure 3F). Whilst presence of this site indicates accessibility of DNA to transcription factors that could activate expression in mesophyll cells, global analysis of all genes strongly and preferentially expressed in bundle sheath strands versus whole leaves indicates that presence/absence of a DHS in one cell type is not sufficient to generate cell specificity (Supplemental Figure 7 & 8). Second, in the next contiguous gene that encodes an additional isoform of MDH also preferentially expressed in mesophyll cells (Figure 3E), a DHS was found in both whole leaf and bundle sheath strands but DGF within this region differed between cell types (Figure 3F). Thus, despite similarity in DNA accessibility, the binding of particular transcription factors varied between cell types. However, once again, genome-wide analysis indicated that alterations to individual DGF were not sufficient to explain cell specific gene expression. For example, only 30 to 40% of all enriched DGF in the bundle sheath were associated with differentially expressed genes (Supplemental Table 8). Lastly, in the third gene in this region, which encodes a NAC domain transcription factor preferentially expressed in bundle sheath strands (Figure 3E), differentially enriched DGF were associated both with regions of the gene that have similar DHS in each cell type, but also a region lacking a DHS in whole leaves compared with bundle sheath strands (Figure 3E). These three classes of alteration to transcription factor accessibility and binding were detectable in genes encoding core components of the C_4_ cycle in all three species (Supplemental Figure 9-11). Overall, we conclude that differences in transcription factor binding between cells of C_4_ leaves is associated with both DNA accessibility defined by broad DHS, as well as fine-scale alterations to transcription factor binding defined by DGF. Moreover, bundle sheath strands possessed a distinct regulatory landscape compared with the whole leaf, and in genes encoding enzymes of the C_4_ pathway multiple transcription factor binding sites differed between bundle sheath and whole leaf samples. This finding implies that cell specific gene expression in C_4_ leaves is mediated by combinatorial effects derived from alterations to gene accessibility as defined by DHS as well as changes to binding of multiple transcription within these regions.

### DNA motifs associated with cell specific expression

To provide an overview of the transcription factors most likely associated with DGF ChIP-SEQ data from maize (Figure 2B) together with motifs from JASPAR plants (Khan et al., 2018) and an additional 529 transcription factor motifs validated in Arabidopsis (O’Malley et al., 2016) were used to annotate the DGF from *Z. mays, S. bicolor, S. italica* and *B. distachyon* (Figure 4A). To increase the number of annotated DGF *de novo* prediction was used to identify sequences over-represented in DGF compared with those across the whole genome. This resulted in an additional 524 motifs being annotated (Figure 4A), but in fact all of these were previously detected after *de novo* prediction from DNaseI-SEQ of rice (Zhang et al., 2012b). As would be expected from *bona fide* transcription factor binding, inspection of these motifs predicted *de novo* demonstrated clear strand bias in DNaseI cuts (Figure 4B). By combining previously known motifs and those predicted *de novo*, the percentage of DGF that could be annotated in each species increased from about 60% to more than 75% (Figure 4C, Supplemental Table 9).

**Figure 4:**
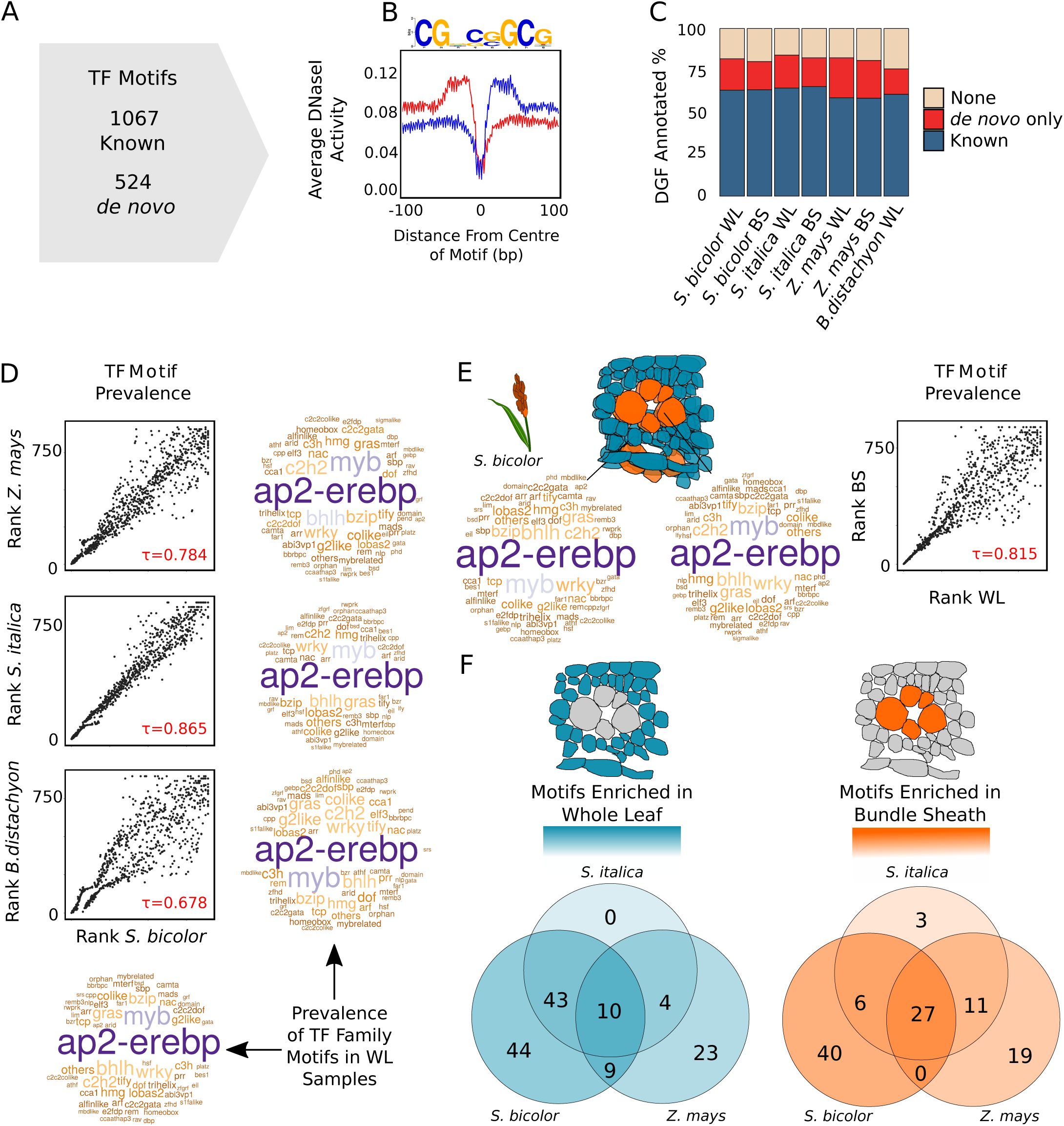
Cistromes associated with cell specific gene expression in C_4_ grasses. (A) Number of previously reported motifs as well as those defined *de novo* in the grasses. (B) Density plots depicting average DNaseI activity on positive (red) and negative (blue) strands centred around a *de novo* motif. (C) Bar chart depicting percentage of DGF annotated with known or *de novo* motifs. (D) Comparison of transcription factor motif prevalence in Whole Leaf (WL) samples from *S. italica, Z. mays, B. distachyon* compared with *S. bicolor*. Word clouds depict frequency of motifs associated with transcription factor families, with larger names more abundant. Scatter plots compare frequency of transcription factor motifs within DGF, ranked from low (most abundant) to high (least abundant). Correlation between samples is indicated as Kendall’s Tau coefficient (τ). (E) Comparison of transcription factor motif prevalence in BS enriched and WL enriched DGF from *S. bicolor*, as in (D), word clouds depict frequency of motifs associated with transcription factor families and plots compare frequency of transcription factor motifs within DGF ranked from low to high. Similarly scatter plots compare transcription factor motif prevalence in BS enriched and whole leaf enriched DGF from *S. bicolor*. (F) Venn diagram showing enriched motifs for each cell type in all three C_4_ species.

To define the most common sequences bound by transcription factors in mature leaves undertaking C_3_ and C_4_ photosynthesis and to investigate whether C_4_ photosynthesis is controlled by an increase in binding of sets of transcription factors, individual motifs were ranked by frequency and the Kendall rank correlation coefficient used to compare species (Figure 4D). In both C_3_ and C_4_ species, the most prevalent transcription factor binding motifs were associated with AP2-EREBP and MYB transcription factor families (p-value < 2.2^-16^; Figure 4D). Next, to identify regulatory factors associated with gene expression in the C_4_ bundle sheath, transcription factor motifs located in DGF enriched in either the bundle sheath or in whole leaf samples of *S. bicolor* were identified (Figure 4E). There was little difference in the ranking of the most commonly used motifs between these cell types (Kendall’s tau=0.815; p-value < 2.2^-16^), indicating cell-specificity is not associated with large-scale changes in the abundance of many transcription factor families (Figure 4E). After performing hypergeometric tests for enrichment of individual motifs in differentially occupied DGF we found 133 and 106 motifs enriched in whole leaves and bundle sheath strands respectively (p < 0.001). Of these 239 motifs, 37 were enriched in all C_4_ species with 10 and 27 enriched in the bundle sheath and whole leaf respectively (Figure 4F, Supplemental Table 10) 66 were only enriched in bundle sheath strands and 91 to whole leaf tissue (Supplemental Table 11). Some of these conserved and cell specific motifs have been previously described to have a relevant role in photosynthesis. For instance, in whole leaves of maize and *Setaria*, we found significant enrichment of the bHLH129 motif (Supplemental Table 11) that has been proposed to act as a negative regulator of NADP-ME (Borba et al., 2018).

### Multiple genes encoding enzymes of the C_4_ and Calvin-Benson-Bassham cycles share the same occupied *cis*-elements

To investigate whether genes involved in the C_4_ phenotype are co-regulated, we compared the number of instances where the same motifs were bound in multiple C_4_, Calvin-Benson-Bassham and C_2_ cycle genes (Supplemental Table 12). While no single *cis*-element was found in all genes that are preferentially expressed in mesophyll or bundle-sheath cells, the number of genes possessing the same occupied motif ranged from nine in *S. bicolor* and *S. italica* to four in *S. bicolor* and *Z. mays* whole leaves respectively (Supplemental Table 9 & 12). These data support a model where the combinatorial action of multiple transcription factors controls groups of C_4_ genes to produce the gene expression patterns required for C_4_ photosynthesis.

We next performed comparative analysis of motifs bound by transcription factors to determine whether the set of *cis*-elements found in C_4_ genes of each species were common, or whether C_4_ genes are regulated differently in each species. In pairwise comparisons, DGF fell into three categories: conserved and occupied by a transcription factor, conserved but only occupied in one species, and not conserved (Figure 5A). Only a small percentage of DGF were both conserved in sequence and bound by transcription factors (Figure 5B, Supplemental Table 13). Consistent with this, the majority of C_4_ gene orthologs did not share DGF. Due to the lack of DGF saturation in maize, these estimates likely set lower bounds for the extent of conservation. However, in several cases, patterning of C_4_ gene expression correlated with a set of motifs shared across species (Figure 5C). In some cases, these shared *cis*-elements were present in the ancestral C_3_ state. For instance, the *TRANSKETOLASE* (*TKL*) gene contains several conserved DGF that are present in the bundle sheath of the C_4_ species but also in whole leaves of C_3_ *Brachypodium* (Figure 5). This finding is consistent with the notion that C_4_ photosynthesis makes use of existing regulatory architecture found in C_3_ plants. Nevertheless, overall, these data also indicate that the majority of C_4_ gene expression appears to be associated with species-specific regulatory networks.

**Figure 5:**
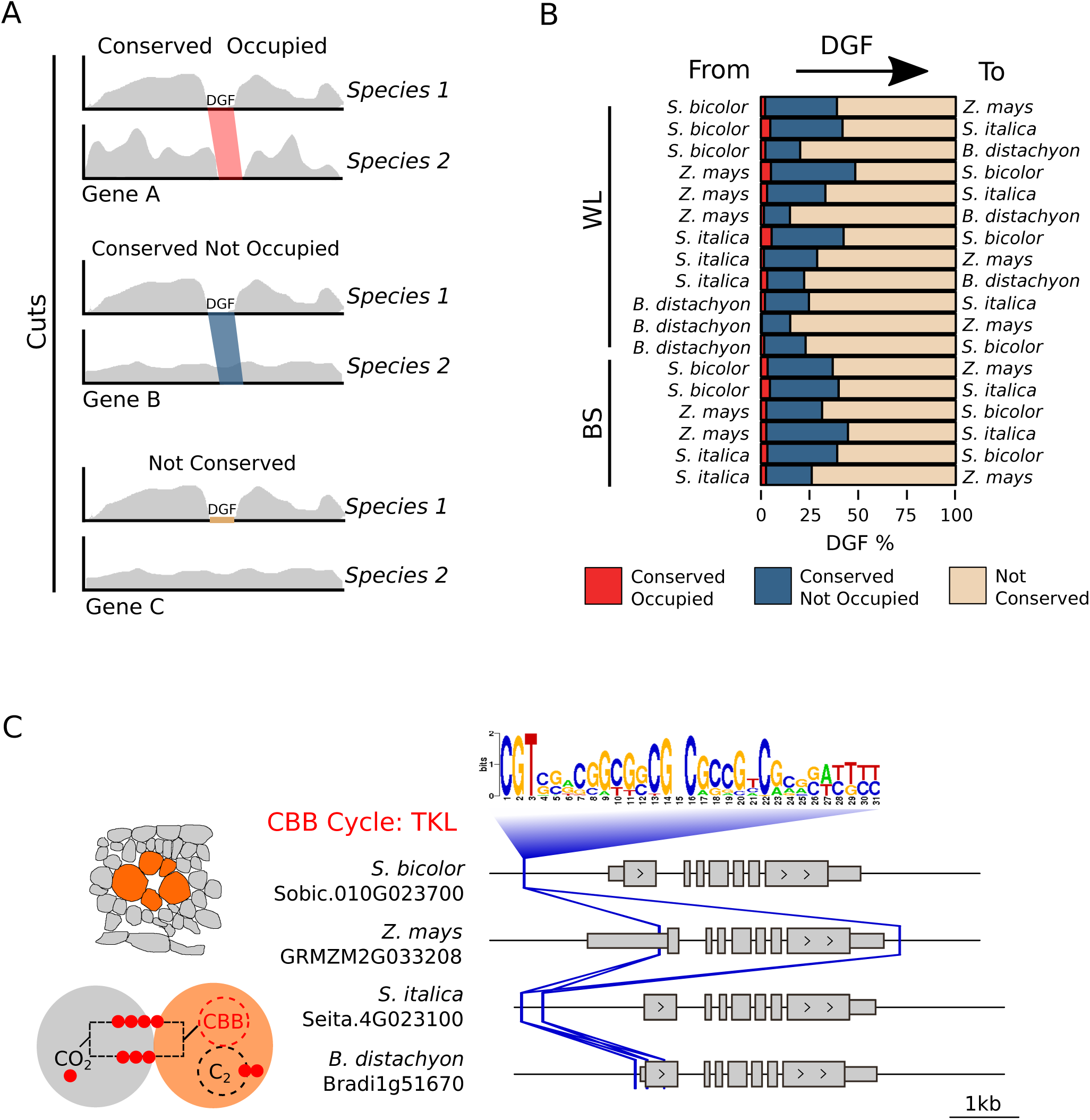
*Cis*-elements show high rates of turnover and mobility in grasses. (A) Scenarios for DGF conservation between species. Reads derived from DNaseI cuts are depicted in grey, DGF that are both conserved and occupied between species by red, and DGF that are conserved but unoccupied by blue shading. (B) Bar-plot representing pairwise comparisons of DGF occupancy. (C) Schematic depicting the position of a transcription factor motif consistently associated with the bundle sheath enriched *TRANSKETOLASE* (*TKL)* gene in *S. bicolor, Z. mays, S. italica* and C_3_ *B. distachyon*. The position of motifs conserved between orthologous genes is depicted by solid lines and orange) and varies between species.

### Hyper-conserved *cis*-regulators of C_4_ genes

To investigate the extent to which transcription factor binding sites associated with C_4_ genes within a C_4_ lineage are conserved, genes encoding the core C_4_ cycle were compared in *S. bicolor* and *Z. mays* (Figure 6A, Supplemental Table 14). 27 genes associated with the C_4_ and Calvin-Benson-Bassham Cycles contained a total of 379 DGF. Although many of these transcription factor footprints were conserved in sequence within orthologous genes, only nine were both conserved and bound by a transcription factor (Figure 6A). Again, due to the lack of DGF saturation in maize, these data likely represent minimum estimates of conservation.

**Figure 6:**
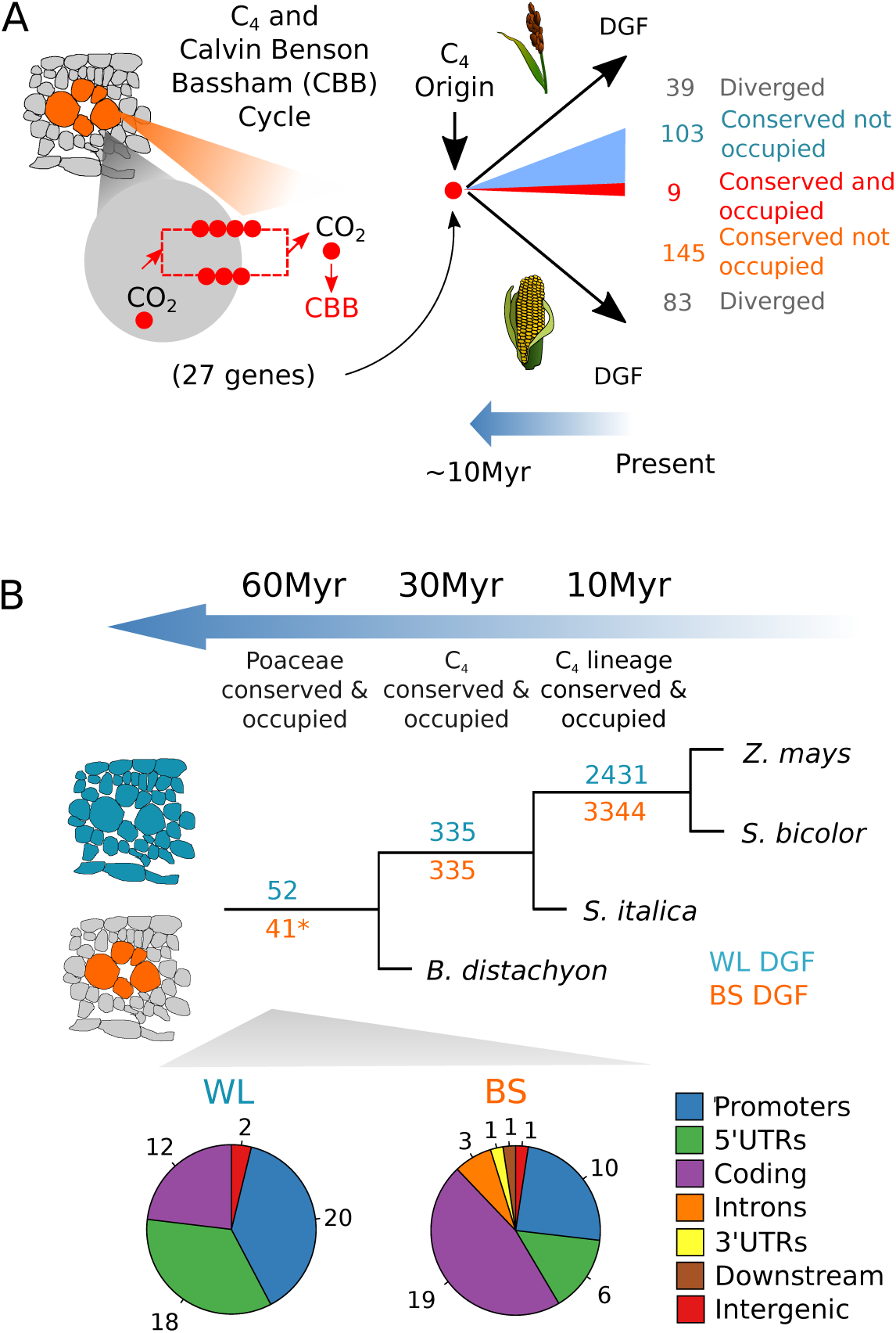
Hyper-conserved *cis*-elements in grasses recruited into C_4_ photosynthesis. (A) Conservation of regulation in C_4_ and Calvin Benson Bassham cycle genes following the divergence of *Z. mays* and *S. bicolor*. The number of carbon atoms (red dots) and metabolite flow (red dashed line) between mesophyll (grey) and bundle sheath (orange) cells are illustrated along with the degree of conservation of DGF associated with BS strands. (B) Conservation of DGF occupancy in grasses across evolutionary time. Results are depicted for whole leaf (WL - blue) and bundle sheath (BS - orange) DGF. The asterisk indicates 41 DGF that are conserved in the BS of the C_4_ species but are also found in whole leaves of *B. distachyon*). Pie-charts display the distribution of conserved and occupied DGF for whole leaf and BS strands. Promoters are defined as sequence up to 2000 base pairs (bp) upstream of the transcriptional start site, downstream represent regions 1000 downstream the transcription termination site while intergenic represent > 1000 bp downstream the transcription termination site until the next promoter region.

Genome-wide, the number of DGF that were conserved in sequence and bound by a transcription factor decayed in a non-linear manner with phylogenetic distance (Figure 6B, Supplemental Table 15). For example, *Z. mays* and *S. bicolor* shared 5,775 DGF that were both conserved and occupied. *S. italica* shared only 670 DGF with *Z. mays* and *S. bicolor* (Figure 6B). Finally, comparison of these C_4_ grasses with C_3_ *B. distachyon* yielded 93 DGF that have been conserved over >60Myr of evolution. Because nuclei from *B. distachyon* were sampled later in the photoperiod than those from the C_4_ grasses, and DGF may well vary over the diel cycle, it is possible that this is an underestimate of DGF conservation. However, 41 of these highly conserved DGF were present in whole leaf samples of the C_3_ species, but in the C_4_ species were restricted to the bundle sheath (Figure 6B). Gene Ontology analysis did not detect enrichment of any specific terms for hyper-conserved DGF associated with the bundle sheath, but for whole leaves detected over-representation of “cell component” categories such as membrane bound organelles and the nucleus (Supplemental Table 16). In whole leaves, this set of ancient and highly conserved DGF were located predominantly in 5’ UTRs and coding sequences, but in bundle sheath strands over fifty percent of these hyper-conserved DGF were in coding sequences (Figure 6B). Overall, these data indicate that certain duons are highly conserved across deep evolutionary time. The frequent use of hyper-conserved duons in the bundle sheath implies that this cell type uses an ancient and highly conserved regulatory code.

## Discussion

### Genome-wide transcription factor binding in grasses

The dataset provides insight into the regulation of gene expression in cereals in general, and to C_4_ photosynthesis in particular. Consistent with previous analysis ranging from *A. thaliana* (Sullivan et al., 2014) to metazoans (Natarajan et al., 2012; Stergachis et al., 2013, 2014), the majority of DGF detected in the four grasses were centred around annotated transcription start sites. However, in these cereals it is noteworthy that transcription factor binding was prevalent in genic sequence. Whilst we cannot rule out the possibility that this distribution is in some way related to the methodology used in this study, there is evidence that the exact distribution of transcription factor binding appears to be species specific. For example, whilst in *A. thaliana* DNaseI-SEQ revealed enrichment of DHS in sequence ∼400 base pairs upstream of transcription start sites as well as 5’ UTRs (Sullivan et al., 2014) and ATAC-SEQ of *A. thaliana, Medicago truncatula* and *Oryza sativa* detected most transposase hypersensitive sites upstream of genes, in *Solanum lycopersicum* more were present in introns and exons than upstream of annotated transcription start sites (Maher et al., 2017). The prevalence of transcription factor binding to coding sequences is relevant to approaches used to generate transgenic plants and test gene function and regulation. First, consistent with the prevalence of DGF downstream of the annotated transcription start sites that we detected, it is noteworthy that during cereal transformation, exon and intron sequences are frequently used to achieve stable expression of transgenes (Maas et al., 1991; Cornejo et al., 1993; Jeon et al., 2000). It is possible that this strategy is required in grasses because of the high proportion of transcription factor binding downstream of annotated transcription start sites. These transcription factor binding sites in coding sequence also have implications for synthetic biology. Although technologies such as type-IIS restriction endonuclease cloning methods allow high-throughput testing of many transgenes, they rely on sequence domestication. Whilst routinely this would maintain amino acid sequence, without analysis of transcription factor binding sites it could mutate motifs bound by transcription factors and lead to unintended modifications to gene expression.

### The transcription factor landscape underpinning gene expression in specific tissues

The finding that so few transcription factor binding sites were shared between bundle sheath tissue and whole leaves of *S. bicolor, Z. mays* and *S. italica* argues for the need to isolate these cells when attempting to understand the control of gene expression. Although separating bundle sheath strands from C_4_ leaves is relatively trivial (Covshoff et al., 2013; Furbank et al., 1985; Leegood, 1985) this is not the case for C_3_ leaves. Approaches in which nuclei from specific cell-types are labelled with an exogenous tag (Deal and Henikoff, 2011) now allow their transcription factor landscapes to be defined. The application of DNaseI-SEQ to specific cell types has recently been used in roots (Maher et al., 2017) and so in the future, this approach of both C_3_ and C_4_ leaves should provide insight into how the extent to which gene regulatory networks have been re-wired during the evolution of the complex C_4_ trait.

Given the central importance of cellular compartmentation to C_4_ photosynthesis, there have been significant efforts to identify *cis*-elements that restrict gene expression to either mesophyll or bundle sheath cells of C_4_ leaves (Hibberd and Covshoff, 2010; Sheen, 1999; Wang et al., 2014). As previous studies of C_4_ gene regulation have focused on individual genes and have been performed in various species, it has not been possible to obtain a coherent picture of regulation of the C_4_ pathway and along with many other systems, initial analysis focussed on regulatory elements located in promoters of C_4_ genes (Sheen, 1999). However, it has become increasingly apparent that the patterning of gene expression between cells in the C_4_ leaf can be mediated by elements in various parts of a gene. In addition to promoter elements (Sheen, 1999; Gowik et al., 2004), this includes untranslated regions (Kajala et al., 2011; Patel et al., 2004; Viret et al., 1994; Williams et al., 2016; Xu et al., 2001) and coding sequences (Brown et al., 2011; Reyna-Llorens et al., 2018). By providing data on *in vivo* transcription factor occupancy for the complete C_4_ pathway in three C_4_ grasses, the data presented here allow broad comparisons and provide several insights into regulatory networks controlling C_4_ genes.

The DNaseI dataset indicates that cell specific gene expression in C_4_ leaves is not strongly correlated with changes to large-scale accessibility of DNA as defined by DHS. This implies that modifications to transcription factor accessibility around any one gene does not impact on its expression between tissues in the leaf. Rather, as only 8-24% of transcription factor binding sites detected in the bundle sheath were also found in whole leaves, the data strongly implicate complex modifications to patterns of transcription factor binding in controlling gene expression between cell types. These findings are consistent with analogous analysis in roots where genes with clear spatial patterns of expression are bound by multiple transcription factors (Sparks et al., 2017) and highly combinatorial interactions between multiple activators and repressors tune the output (de Lucas et al., 2016).

The data also provide insight into *cis*-elements that underpin the C_4_ phenotype. No single c*is*-element was found in all genes preferentially expressed in either mesophyll or bundle sheath cells of one species. This finding is consistent with analysis of yeast where the output of genetic circuits can be maintained despite rapid turnover of *cis-*regulatory mechanisms underpinning them (Tsong et al., 2006). However, we did detect small numbers of C_4_ genes that shared common transcription factor footprints (Figure 5, Supplemental Table 14 & 15), which is consistent with previous analysis that identified shared *cis*-elements in *PPDK* and *CA,* or *NAD-ME1* and *NAD-ME2* in C_4_ *Gynandropsis gynandra* (Williams et al., 2016; Reyna-Llorens et al., 2018). Interspecific comparisons further underlined the high rate of divergence in the *cis*-regulatory logic used to control C_4_ genes. For example, although we detected highly similar transcription factor footprints in the *OMT1* and *TKL* genes of the three C_4_ species we assessed, this was not apparent for any other C_4_ genes. As a result of the apparent rapid rate of evolution in *cis*-regulatory architecture in these C_4_ species, attempts to engineer C_4_ photosynthesis into C_3_ crops to increase yield (Hibberd et al., 2008) may benefit from using pre-existing regulatory mechanisms controlling mesophyll or bundle sheath expression in ancestral C_3_ species.

### Characteristics of the transcription factor binding in the ancestral C_3_ state that have impacted on evolution of the C_4_ pathway

Comparison of transcription factor binding in the C_3_ grass *B. distachyon* with three C_4_ species provides insight into mechanisms associated with the evolution of C_4_ photosynthesis. For all four grasses, irrespective of whether they used C_3_ or C_4_ photosynthesis, the most abundant DNA motifs bound by transcription factors were similar. Thus, motifs recognised by the AP2-EREBP and MYB classes of transcription factor were most commonly bound across each genome. This indicates that during the evolution of C_4_ photosynthesis, there has been relatively little alteration to the most abundant classes of transcription factors that bind DNA.

The repeated evolution of the C_4_ pathway has frequently been associated with convergent evolution (Sage, 2004; Sage et al., 2012). However, parallel alterations to amino acid and nucleotide sequence that allow altered kinetics of the C_4_ enzymes (Christin et al., 2014, 2007) and patterning of C_4_ gene expression (Brown et al., 2011) respectively have also been reported. The genome-wide analysis of transcription factor binding reported here indicates that only a small proportion of the C_4_ cistrome is associated with parallel evolution. These estimates regarding conservation between C_4_ and C_3_ species may represent underestimates as whilst nuclei where all sampled in the light, those from *B. distachyon* were sampled later in the photoperiod. Moreover, when orthologous genes were compared between the four grasses assessed here, the majority of transcription factor binding sites were not conserved, and of the DGF that were conserved, position within orthologous genes varied. This indicates that C_4_ photosynthesis in grasses is tolerant to a rapid turnover of the *cis*-code, and that when motifs are conserved in sequence, their position and frequency within a gene can vary. It therefore appears that the cell-specific accumulation patterns of C_4_ proteins can be maintained despite considerable modifications to the cistrome of C_4_ leaves. It was also the case that some conserved motifs bound by transcription factors in the C_4_ species were present in *B. distachyon*, which uses the ancestral C_3_ pathway. Previous work has shown that *cis*-elements used in C_4_ photosynthesis can be found in gene orthologs from C_3_ species (Williams et al., 2016; Reyna-Llorens et al., 2018). However, these previous studies identified *cis*-elements that were conserved in both sequence and position. As it is now clear that such conserved motifs are mobile within a gene, it seems likely that many more examples of ancient *cis*-elements important in C_4_ photosynthesis will be found in C_3_ plants.

Although we were able to detect a small number of transcription factor binding sites that were conserved and occupied in all four species sampled, these ancient hyper-conserved motifs appear to have played a role in the evolution of C_4_ photosynthesis. Interestingly, a large proportion of these motifs bound by transcription factors were found in coding sequences, and this bias was particularly noticeable in bundle sheath cells. Due to the amino acid code, the rate of mutation of coding sequence compared with the genome is restricted. If such regions have a longer half-life than transcription factor binding sites in other regions of the genome, then they may represent an excellent source of raw material for the repeated evolution of complex traits (Martin and Orgogozo, 2013). Our data documenting the frequent use of hyper-conserved DGF in the C_4_ bundle sheath implies that this tissue may use an ancient and highly conserved regulatory code. It appears that during the evolution of the C_4_ pathway, which relies on heavy use of the bundle sheath, this ancient code has been co-opted to control photosynthesis gene expression.

In summary, the data provide a transcription factor binding atlas for leaves of grasses using either C_3_ or C_4_ photosynthesis. Whilst we did not achieve DGF saturation in maize, commonalities between the four species were apparent. Sequences bound by transcription factors were found within genes as well as promoter regions, and many of these motifs represent duons. In terms of the regulation of tissue specific gene expression, whilst bundle sheath strands and whole C_4_ leaves shared considerable similarity in regions of DNA accessible to transcription factors, the short sequences actually bound by transcription factors varied dramatically. We identified a small number of transcription factor motifs that were conserved in these species. The data also provide insight into the regulatory architecture associated with C_4_ photosynthesis more specifically. Whilst we found some evidence that multiple genes important for C_4_ photosynthesis share common *cis*-elements bound by transcription factors, this was not widespread. This may well relate to the relatively rapid turnover in the *cis*-code, and so it is possible that transcription factors interacting with these motifs are more conserved. Analysis of transcription factor footprints in specific cell types from leaves of C_3_ grasses should in the future provide insight into the extent to which gene regulatory networks have altered during the transition from C_3_ to C_4_ photosynthesis.

## Methods

### Growth conditions and isolation of nuclei

*S. bicolor, S. italica* and *Z. mays* were grown under controlled conditions at the Plant Growth Facilities of the Department of Plant Sciences at the University of Cambridge in a chamber set to 12 h/12 h light/dark; 28 °C light/20 °C dark; 400 µmol m^-2^ s^-1^ photon flux density, 60% humidity. For germination, *S. bicolor* and *Z. mays* seeds were imbibed in H_2_O for 48 h, *S. italica* seeds were incubated on wet filter paper at 30 °C overnight in the dark. Z*. mays, S. bicolor* and *S. italica* were grown on 3:1 (v/v) M3 compost to medium vermiculite mixture, with a thin covering of soil. Seedlings were hand-watered.

*B. distachyon* plants were grown in a separate growth facility under controlled conditions optimised for its growth at the Sainsbury Laboratory Cambridge University, first under short day conditions 14 h/10 h, light/dark for 2 weeks and then shifted to long day 20 h/4 h, light/dark, for 1 week and harvested at ZT20. Temperature was set at 20 °C, humidity 65% and light intensity 350 µmol m^-2^ s^-1^. All tissue was harvested from August to October 2015.

To isolate nuclei from *S. bicolor, Z. mays* and *S. italica* mature third and fourth leaves with a fully developed ligule were harvested 4-6 h into the light cycle 18 days after germination. Bundle sheath cells were mechanically isolated as described previously (Markelz et al., 2003). At least 3 g of tissue was used for each extraction. Nuclei were isolated using a sucrose gradient adapted and yield quantified using a haemocytometer. For *B. distachyon* plants were flash frozen and material pulverised in a coffee grinder. 3 g of plant material was added to 45 ml NIB buffer (10mM Tris-HCl, 0.2M sucrose, 0.01% (v/v) Triton X-100, pH 5.3 containing protease inhibitors (Sigma-Aldrich)) and incubated at 4°C on a rotating wheel for 5 min, afterwards debris was removed by sieving through 2 layers of Miracloth (Millipore) into pre-cooled flasks. Nuclei were spun down 4,000 rpm, 4 °C for 20 min. Plastids were lysed by adding Triton to a final concentration of 0.3% (v/v) and incubated for 15 min on ice. Nuclei were pelleted by centrifugation at 5000 rpm at 4 °C for 15 min. Pellets were washed 3 times with chilled NIB buffer.

### Deproteinized DNA extraction

For isolation of deproteinated DNA from *S. bicolor*, *Z. mays*, *B. distachyon* and *S. italica* mature third and fourth leaves with a fully developed ligule were harvested 4 h into the light cycle, 18 days after germination. 100 mg of tissue was used for each extraction. Deproteinated DNA was extracted using a QIAGEN DNeasy Plant Mini Kit (QIAGEN, UK) according to the manufacturer’s instructions.

### DNaseI digestion, sequencing and library preparation

To obtain sufficient DNA each biological replicate consisted of leaves from tens of individuals and to conform to standards set by the Human Genome project at least two biological replicates were sequenced for each sample. 2 x 10^8^ of freshly extracted nuclei were re-suspended at 4 °C in digestion buffer (15 mM Tris-HCl, 90 mM NaCl, 60 mM KCl, 6 mM CaCl_2_, 0.5 mM Spermidine, 1 mM EDTA and 0.5 mM EGTA, pH 8.0). DNaseI (Fermentas) at 7.5 U was added to each tube and incubated at 37 °C for 3 min. Digestion was arrested with addition of 1:1 volume of stop buffer (50 mM Tris-HCl, 100 mM NaCl, 0.1% (w/v) SDS, 100 mM EDTA, pH 8.0, 1 mM Spermidine, 0.3 mM Spermine, RNase-A 40 µg/ml) and incubated at 55 °C for 15 min. 50 U of Proteinase K was added and samples incubated at 55 °C for 1 h. DNA was isolated with 25:24:1 Phenol:Chloroform:Isoamyl Alcohol (Ambion) followed by ethanol precipitation. Fragments from 50 to 550 bp were selected using agarose gel electrophoresis. The extracted DNA samples were quantified fluorometrically with a Qubit 3.0 Fluorometer (Life Technologies), and a total of 10 ng of digested DNA (200 pg l-1) was used for library construction.

Initial sample quality control of pre-fragmented DNA was assessed using a Tapestation DNA 1000 High Sensitivity Screen Tape (Agilent, Cheadle UK). Sequencing ready libraries were prepared using the Hyper Prep DNA Library preparation kit (Kapa Biosystems, London UK) selecting fragments from 70-350 bp for optimization (see He et al., 2014) and indexed for pooling using NextFlex DNA barcoded adapters (Bioo Scientific, Austin TX US). In order to reduce bias due to amplification of DNA fragments by the polymerase chain reaction, as recommended by the manufacturers, a low number of cycles (17 cycles) was used. Libraries were quantified using a Tapestation DNA 1000 Screen Tape and by qPCR using an NGS Library Quantification Kit (KAPA Biosystems) on an AriaMx qPCR system (Agilent) and then normalised, pooled, diluted and denatured for sequencing on the NextSeq 500 (Illumina, Chesterford UK). The main library was spiked at 10% with the PhiX control library (Illumina). Sequencing was performed using Illumina NextSeq in the Departments of Biochemistry and Pathology at the University of Cambridge, UK, with 2x75 cycles of sequencing. For the deproteinized DNAse I seq experiments 1 µg of deproteinized DNA was resuspended in 1 ml of digestion buffer (15 mM Tris-HCl, 90 mM NaCl, 60 mM KCl, 6 mM CaCl_2_, 0.5 mM spermidine, 1 mM EDTA and 0.5 mM EGTA, pH 8.0). DNaseI (Fermentas) at 2.5 U was added to each tube and incubated at 37 °C for 2 min. Digestion was arrested with addition of 1:1 volume of stop buffer (50 mM Tris-HCl, 100 mM NaCl, 0.1% (w/v) SDS, 100 mM EDTA, pH 8.0, 1 mM Spermidine, 0.3 mM Spermine, RNase A 40 µg/ml) and incubated at 55 °C for 15 min. 50 U of Proteinase K was added and samples incubated at 55 °C for 1 h. DNA was isolated by mixing with 1 ml 25:24:1 Phenol:Chloroform:Isoamyl Alcohol (Ambion) and spun for 5 min at 13,00 rpm followed by ethanol precipitation of the aqueous phase. Samples were then size-selected (50-400 bp) using agarose gel electrophoresis. The extracted DNA samples were quantified fluorometrically using Qubit 3.0 Fluorometer (Life technologies), and a total of 1 ng of digested DNA was used for library construction. Sequencing ready libraries were prepared using a KAPA Hyper Prep Kit (KAPA Biosystems, London UK) according to the manufacturer’s instructions. In order to reduce bias due to amplification of DNA fragments by the polymerase chain reaction, as recommended by the manufacturers, 17 cycles were used. Quality of the libraries were checked using a Bioanalyzer High Sensitivity DNA Chip (Agilent Technologies). Libraries were quantified by Qubit 3.0 Fluorometer (Life Technologies) and qPCR using an NGS Library Quantification Kit (KAPA Biosystems) and then normalised, pooled, diluted and denatured for paired end sequencing using High Output 150 cycle run (2x 75 bp reads). Sequencing was performed using NextSeq 500 (Illumina, Chesterford UK) in the Sainsbury Laboratory University of Cambridge, UK, with 2x75 cycles of sequencing.

### DNaseI-SEQ Data processing

Genome sequences were downloaded from Phytozome (v10) (Goodstein et al., 2012). The following genome assemblies were used: Bdistachyon_283_assembly_v2.0; Sbicolor_255_v2.0; Sitalica_164_v2; Zmays_284_AGPv3. Due to the lack of guidelines for DNaseI-SEQ experiments in plants we followed the guidelines from the Encyclopaedia of DNA Elements (ENCODE3). Reads were mapped to genomes using bowtie2 (Langmead and Salzberg, 2012) and processed using samtools (Li et al., 2009) to remove those with a MAPQ score <42. DHS were called using MACS2 (Feng et al., 2012) and the final set of peak calls were determined using the irreproducible discovery rate (IDR) (Li and Dewey, 2011), calculated using the script batch_consistency_analysis.R (https://github.com/modENCODE-DCC/Galaxy/blob/master/modENCODE_DCC_tools/idr/batch-consistency-analysis.r). The Irreproducible Discovery Rate framework adapted from the ENCODE 3 pipeline (Marinov et al., 2014; https://sites.google.com/site/anshulkundaje/projects/idr) aims to measure the reproducibility of findings by identifying the point (threshold) in which peaks are no longer consistent across replicates.

### Quality metrics and identification of Digital Genomic Footprints (DGF)

SPOT score (number of a subsample of mapped reads (5M) in DHS/Total number of subsampled, mapped reads (5M) (John et al., 2011)) was calculated using BEDTools (Quinlan and Hall, 2010) to determine the number of mapped reads possessing at least 1 bp overlap with a DHS site. Normalized Strand Cross-correlation coefficient (NSC) and Relative Strand Cross-correlation coefficient (RSC) scores were calculated using SPP (Kharchenko et al., 2008) and PCR bottleneck coefficient (PBC) was calculated using BEDTools. To account for cutting bias associated with the DNaseI enzyme DNaseI-SEQ on naked DNA was performed. These data were used to generate background signal profiles and calculate the footprint log-likelihood ratio for each footprint using the R package MixtureModel (Yardimci et al., 2014) such that those with low log likelihood ratios (FLR <0) were removed. Digital Genomic Footprints (DGF) were identified using Wellington (Piper et al., 2013) and differential DGF were identified using Wellington bootstrap (Piper et al., 2015).

### Data visualisation

DHS and DGF sequences were loaded into and visualized in the Integrative Genomics Viewer (Thorvaldsdóttir et al., 2013) and figures produced in Inkscape, plots were generated with R package ggplot2 (Wickham, 2010) and figures depicting conservation of DGF or motifs between orthologous sequences were generated using genoplotR (Guy et al., 2010). Word clouds were created with the wordcloud R package (Fellows, 2012). TreeView images were produced by processing DGF data using ‘dnase_to_javatreeview.py’ from pyDNAse (Piper et al., 2013, 2015) and loaded into TreeView (Saldanha, 2004). Average cut density plots were generated using the script ‘dnase_average_profile.py’ from pyDNase. Genomic features were annotated and distribution calculated using PAVIS (Huang et al., 2013) and plotted using ggplot2. Circular plots showing the distribution of ChIP-SEQ peaks, DHS and DGF across the maize genome was generated using the R package circlize (Gu, 2014).

### DNAse cutting bias calculations and ChIP-SEQ analysis

After sequencing, the number of DNA 6-mer centred at each DNase cleavage site (between 3^rd^ and 4^th^ base) was counted and normalized by the total number of counts. Next, DNA 6-mer frequencies were normalized by the frequencies of each DNA 6-mer in the genome. The resulting background signal profile was used as input in the FootprintMixture.R package (https://ohlerlab.mdc-berlin.de/software/FootprintMixture_109/) (Supplemental Figure 2).

ChIP-SEQ peaks from 117 transcription factors were obtained from the pre-release maize cistrome data collection (http://www.epigenome.cuhk.edu.hk/C3C4.html). Permutation tests between ChIP peaks and DHS or DGF were performed using regioneR (Gel et al., 2016) using 100 permutations.

### *de novo* motif prediction, motif scanning and enrichment testing

*de novo* motif prediction was performed using findMotifsGenome.pl script from the HOMER suite (Heinz et al., 2010) using digital genomic footprints (DGF) as input together with the reference genome sequence for each species. Motif scanning was performed using FIMO (Grant et al., 2011) with default parameters. To determine overrepresentation of TF family motifs in samples hypergeometric tests were performed using R. The distribution of each motif across different genomic features was obtained for each annotated motif by dividing the number of hits in a particular feature by the total number of hits in the genome.

### Whole genome alignments, pairwise cross mapping of genomic features and variant data processing

To cross map genomic features between species, mapping files were generated according to (http://genomewiki.ucsc.edu/index.php/Whole_genome_alignment_howto) using tools from the UCSC Genome Browser, including trfBig, faToNib, faSize, lavToPsl, faSplit, axtChain, chainNet (Kent et al., 2002) and LASTZ (Harris, 2007). Briefly, whole genome alignment was performed with LASTZ, matching alignments next to each other were chained together using axtChain, sorted with axtSort, then netted together to form larger blocks with chainNet. Genomic features where then mapped between genomes using bnMapper (Denas et al., 2015). For the variant analysis on duons, *Z. mays* variant data (Bukowski et al., 2018) was downloaded from http://cbsusrv04.tc.cornell.edu/users/panzea/download.aspx?filegroupid=16 following instructions. After downloading vcf files were annotated using SnpEff (Cingolani et al, 2012; https://doi.org/10.4161/fly.19695) with the B73_RefGen_v4 genome assembly specifically to allow identification of non-synonymous sites. A custom script was used to identify all four-fold degenerate sites (FFDS) in the *Z. mays* genome. This bed file in turn was used to identify which of the synonymous polymorphic sites were FFDS. Each polymorphic site had its allele frequencies calculated. Putative *Z. mays* duons were identified by intersecting (with bedtools intersect) the final DGF identified with exonic regions. These duons were then used to extract only those exons within which a duon was found. These exons in turn had the duon regions themselves subtracted to leave the exon region except the duon. This provided the surrounding exonic sequences with which to compare to the duons. These two regions were then intersected with the polymorphism data to identify both the number of occurrences and allelic frequencies of polymorphic sites (FFDS and non-synonymous) within both the duons and their surrounding exonic sequences.

### Accession numbers

Methods for DNaseI digestion are on protocols.io (dx.doi.org/10.17504/protocols.io.hdfb23n). Raw sequencing data and processed files are deposited in Gene Expression Omnibus (GSE97369). For full methods, commands and scripts, please see github (https://github.com/hibberd-lab/Burgess-Reyna_llorens-monocot-DNase) and Figshare 10.6084/m9.figshare.7649450.

## Supplemental Legends

**Supplemental Figure 1:** DNaseI digestion of nuclei for sequencing. Representative images of digested samples separated on 2% (w/v) agarose gels by electrophoresis. (A) *S. bicolor* whole leaf (WL); (B) *S. bicolor* Bundle Sheath (BS); (C) *Z. mays* WL; (D) *Z. mays* BS; (E) *B. distachyon* WL; (F) *S. italica* WL; (G) *S. italica* BS. Each gel represents a separate biological replicate, and the units of DNaseI used are illustrated above. Samples selected for sequencing are indicated in red.

**Supplemental Figure 2:** Bias in DNaseI-SEQ cleavage. (A) TreeView diagrams illustrating cut density around individual digital genomic footprint (DGF) predicted from performing DNaseI-SEQ on deproteinated genomic DNA from each species. Each row represents an individual DGF, cuts are coloured according to whether they align to the positive (red) or negative (blue) strand and indicate increased cutting in a 100 bp window on either side of the DGF. (B) Pearson correlation coefficient of DNAse I cleavage bias between *Z. mays*, *S. bicolor*, *S. italica* and *B. distachyon*. (C) Schematic illustrating the process adopted to determine DNaseI cutting bias and then normalize to allow digital genomic footprinting.

**Supplemental Figure 3:** Saturation analysis of footprints. Digital genomic footprints were predicted from subsets (12.5, 25, 50, 75 and 100%) of uniquely mapped reads obtained from DNAseI-SEQ of whole leaf samples in each species.

**Supplemental Figure 4:** Genome wide comparison of DGF and ChIP-SEQ peaks from 117 maize transcription factors. (A) Density plot of DHS, DGF, ChIP-SEQ peaks and intersecting DGF/ChIP-SEQ peaks across the maize genome. The center of the plot shows a word cloud representing transcription factor families in the ChIP-SEQ dataset. (B) Effect of shifting DHS and DGF features from their original position on local z-scores as determined by permutation tests between ChIP-SEQ peaks derived from 117 transcription factors. A total of 100 permutations were performed for each comparison. The sharper peak derived from shifting the DGF indicates a higher sensitivity to position and therefore strong overlap with ChIP-SEQ data. (C) Density plot depicting the distribution of ChIP-SEQ signals per kilobase (kb) from the transcription start site (TSS) of *Z. mays*.

**Supplemental Figure 5:** Density plot depicting the distribution of DGF per kilobase (kb) from the transcription start site (TSS) of *S. bicolor, Z. mays, S. italica* and *B. distachyon* whole leaves.

**Supplemental Figure 6:** Nucleotide proportion of duons and surrounding exons used in the substitution analysis for *Z. mays.* The frequency of each nucleotide was divided by the total length to determine nucleotide proportions across duons as well as surrounding exon sequences.

**Supplemental Figure 7:** Transcript abundance for genes in mesophyll and bundle sheath cells associated with DHS and DGF in *S. bicolor*. (A) Cell preferential gene expression profiles of highly abundant M and BS genes expressed as transcripts per million reads (TPM). (B) Schematic representing DHS, DGF and DE DGF present in whole leaf (blue) and BS (orange) of *S. bicolor*.

**Supplemental Figure 8:** Differential accessibility of broad regulatory regions in *S. bicolor* is not sufficient for cell preferential gene expression. Percentage of differentially detected DHS among BS and M specific genes in *S. bicolor* compared with randomly generated gene samples (n=50).

**Supplemental Figure 9:** Representation of the C_4_ pathway showing differentially accessible DHS, DGF and cell specific DGF between whole leaf (blue) and bundle sheath (orange) samples in *S. bicolor. CA; Carbonic Anhydrase, PEPC; Phosphoenolpyruvate carboxylase, PPDK; Pyruvate, orthophosphate dikinase, MDH; Malate dehydrogenase, NADP-ME; NADP-dependent malic enzyme, RBCS1A; Ribulose bisphosphate carboxylase small subunit1A*, OAA; Oxaloacetate, Mal; Malate, PEP; Phosphoenolpyruvate, Pyr; Pyruvate, Asp; Aspartate.

**Supplemental Figure 10:** Representation of the C_4_ pathway showing differentially accessible DHS, DGF and cell specific DGF between whole leaf (blue) and bundle sheath (orange) samples in *S. italica. CA; Carbonic Anhydrase, PEPC; Phosphoenolpyruvate carboxylase, PPDK; Pyruvate, orthophosphate dikinase, MDH; Malate dehydrogenase, NADP-ME; NADP-dependent malic enzyme, RBCS1A; Ribulose bisphosphate carboxylase small subunit1A*, OAA; Oxaloacetate, Mal; Malate, PEP; Phosphoenolpyruvate, Pyr; Pyruvate, Asp; Aspartate.

**Supplemental Figure 11:** Representation of the C_4_ pathway showing differentially accessible DHS, DGF and cell specific DGF between whole leaf (blue) and bundle sheath

(orange) samples in *Z. mays. CA; Carbonic Anhydrase, PEPC; Phosphoenolpyruvate carboxylase, PPDK; Pyruvate, orthophosphate dikinase, MDH; Malate dehydrogenase, NADP-ME; NADP-dependent malic enzyme, RBCS1A; Ribulose bisphosphate carboxylase small subunit1A*, OAA; Oxaloacetate, Mal; Malate, PEP; Phosphoenolpyruvate, Pyr; Pyruvate, Asp; Aspartate.

**Supplemental Table 1:** Summary of DNAseI-SEQ Quality Metrics. Values include the number of uniquely mapped reads with MAPQ scores >= 42 (NMAP), PCR bottleneck Coefficient (PBC), Normalized Strand Cross-correlation Coefficient (NSC), Relative Strand Cross-correlation Coefficient (RSC) the optimal number of peaks calculated by IDR method (PEAKS), and the Signal Portion Of Tags (SPOT) for whole leaf and bundle sheath samples from *B. distachyon, S. italica, S. bicolor* and *Z. mays*.

**Supplemental Table 2:** Summary Statistics for Genomic Features Identified in Whole Leaf Samples. Including information about the number of DHS and DGF identified per sample, and the number of genes that could be annotated with at least one genomic feature for *B. distachyon, S. italica, S. bicolor* and *Z. mays*.

**Supplemental Table 3:** DNAseI cutting bias calculation summary for whole leaf and bundle sheath data.

**Supplemental Table 4:** Transcription factors included in the ChIP-SEQ data analysis.

**Supplemental Table 5:** Summary statistics of DNAseI-SEQ analysis of Bundle Sheath samples

**Supplemental Table 6:** Summary Statistics of Overlap between DHS in Whole Leaf and Bundle Sheath Samples.

**Supplemental Table 7:** Summary Statistics of Overlap between DGF in Whole Leaf and Bundle Sheath Samples.

**Supplemental Table 8:** Summary Statistics for Differential Digital Genomic Footprint Calling. Including the total number of differential DGF (DE DGF) and the number of DE DGF in DE genes for *S. italica, S. bicolor* and *Z. mays*.

**Supplemental Table 9:** Motifs mapped to genes of the C_4_, CBB and C_2_ cycles in *Z. mays, S. bicolor, S. italica* for whole leaf and bundle sheath samples and in *B. distachyon* for whole leaf samples

**Supplemental Table 10:** Hypergeometric tests for enrichment of individual motifs in *Z. mays, S. bicolor, S. italica* for whole leaf and bundle sheath samples.

**Supplemental Table 11:** Hypergeometric tests for enrichment of cell specific individual motifs in *Z. mays, S. bicolor, S. italica* for whole leaf and bundle sheath samples.

**Supplemental Table 12:** Number of genes in C_4_, CBB and C_2_ cycles annotated with a given motif in *Z. mays, S. italica, S. bicolor* and *B. distachyon*.

**Supplemental Table 12:** Statistics for Cross Mapping of genomic features between *S. bicolor, S. italica, Z. mays* and *B. distachyon*.

**Supplemental Table 13:** DGF conserved and occupied in *Z. mays, S. bicolor, S. italica* for whole leaf and bundle sheath samples and in *B. distachyon* for whole leaf samples.

**Supplemental Table 14:** DGF in C_4_ genes that are conserved between *Z. mays* and *S. bicolor*.

**Supplemental Table 15:** DGF conserved in all four species.

**Supplemental Table 16:** Gene Ontology analysis on hyper-conserved DGF in whole leaf samples of *S. italica, S. bicolor, Z. mays* and *B. distachyon*.

## Acknowledgements

SJB was supported by the 3to4 grant from the EU and BB/I002243 from the BBSRC, IRL by CONACyT and BBSRC grant BB/L014130, SRS and PS by an Advanced ERC grant 694733 Revolution to JMH, and KJ by a Gatsby Career Development Fellowship. We would like to acknowledge Aslihan Karabacak for support in implementing the FootprintMixture package.

## Contributions

SJB, I-RL and JMH conceptualised the experiments. SJB and I-RL grew and harvested nuclei from *S. bicolor, S. italica* and *Z. mays.* KJ provided the nuclei from *B. distachyon*. SJB and I-RL performed DNase I experiments and data analysis. PS extracted nuclei and performed DNAseI experiments on naked DNA, SRS undertook the variant analysis. SJB, I-RL, SRS and JMH wrote the manuscript and prepared the figures.

