## Supplementary material for "Genome-wide transcription factor binding in leaves from C_3_ and C_4_ grasses": S Figures

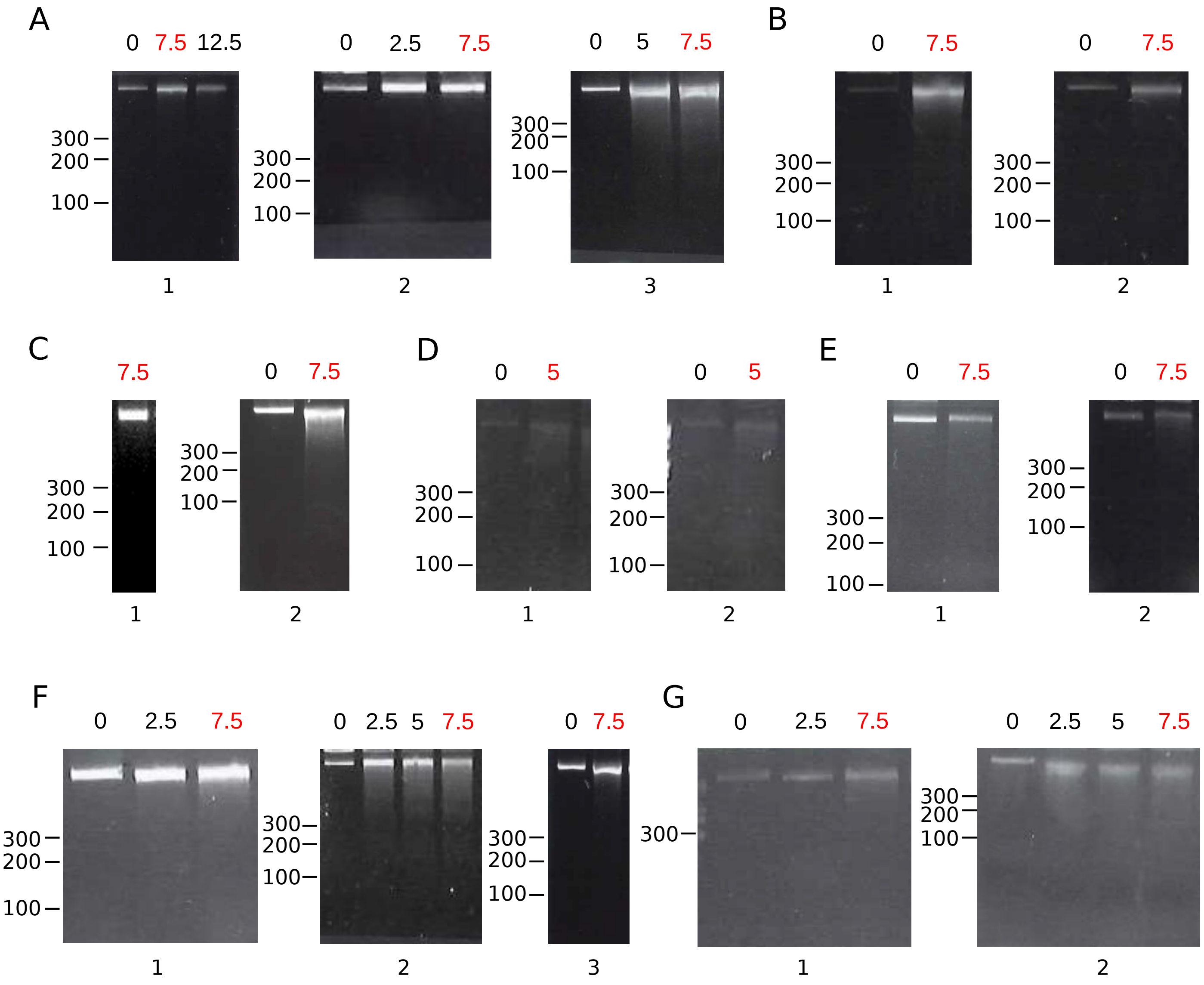


Supplemental Figure 1

A)

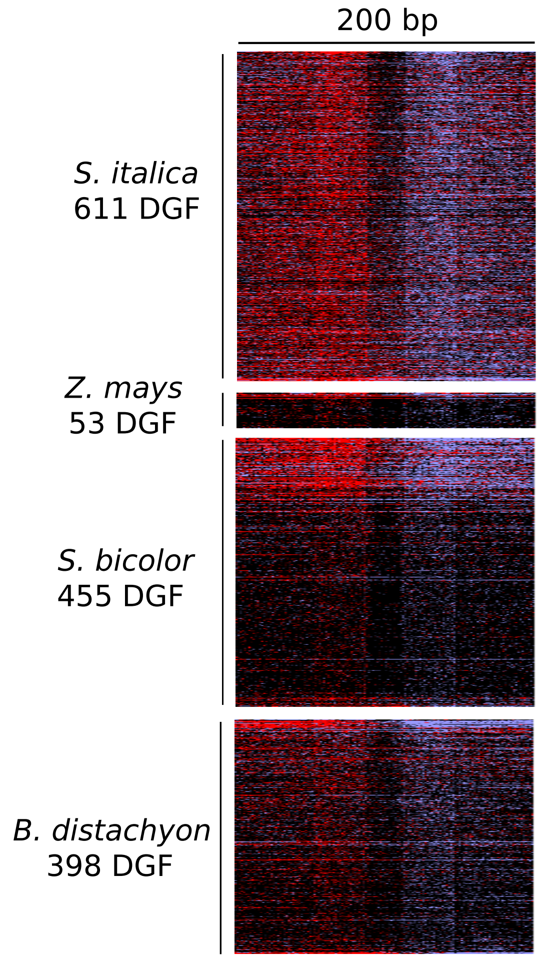


B)

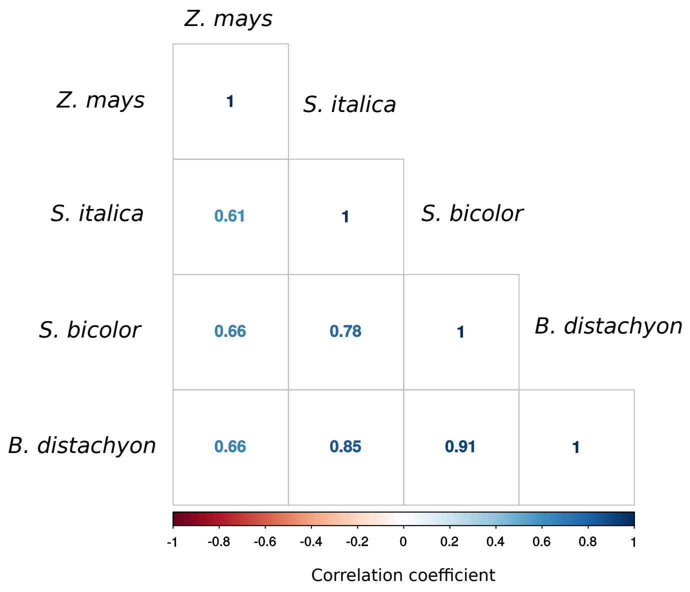


C)

Deproteinized genomic DNA extraction from 4 Monocot Species

↓

DNase-I seq

↓

Naked DNA seq summary

| Species | Tissue | NMAP MAPQ = 42 |
|---|---|---|
| *S. italica* | WL | 18715484 |
| *B. distachyon* | WL | 3935682 |
| *S. bicolor* | WL | 1770650 |
| *Z. mays* | WL | 548194 |

↓

Bias is calculated for each Species by dividing frequency of cuts for each 6-mer by background frequency

↓

| 6-mer | 6-mer freq in the genome | DNase I cut frequency | Cutting bias |
|---|---|---|---|
| AAAAAA | 0.05 | 0.01 | 2 |
| AAAAAT | 0.06 | 0.05 | 0.83 |
| AAAAAC | 0.05 | 0.04 | 2 |
| . | . | . | . |
| . | . | . | . |
| . | . | . | . |
| TTTTTT | n | n | n |

↓

FootPrintMixture.R incorporates bias file to correct DGFs for DNase-I cutting bias

↓

DGF with Footprint Likelihood Ratios (FLR) <0 are filtered out

↓

Digital Genomic Footprinting after filtering

| Sample | DGF | DGF removed after correction | % removed |
|---|---|---|---|
| Sb WL | 142108 | 29813 | 22 |
| Si WL | 120468 | 19994 | 17 |
| Zm WL | 79787 | 23268 | 30 |
| Bd WL | 191046 | 30129 | 16 |
| Sb BS | 128643 | 32968 | 26 |
| Si BS | 158702 | 24042 | 15 |
| Zm BS | 164918 | 23071 | 14 |

Supplemental Figure 2

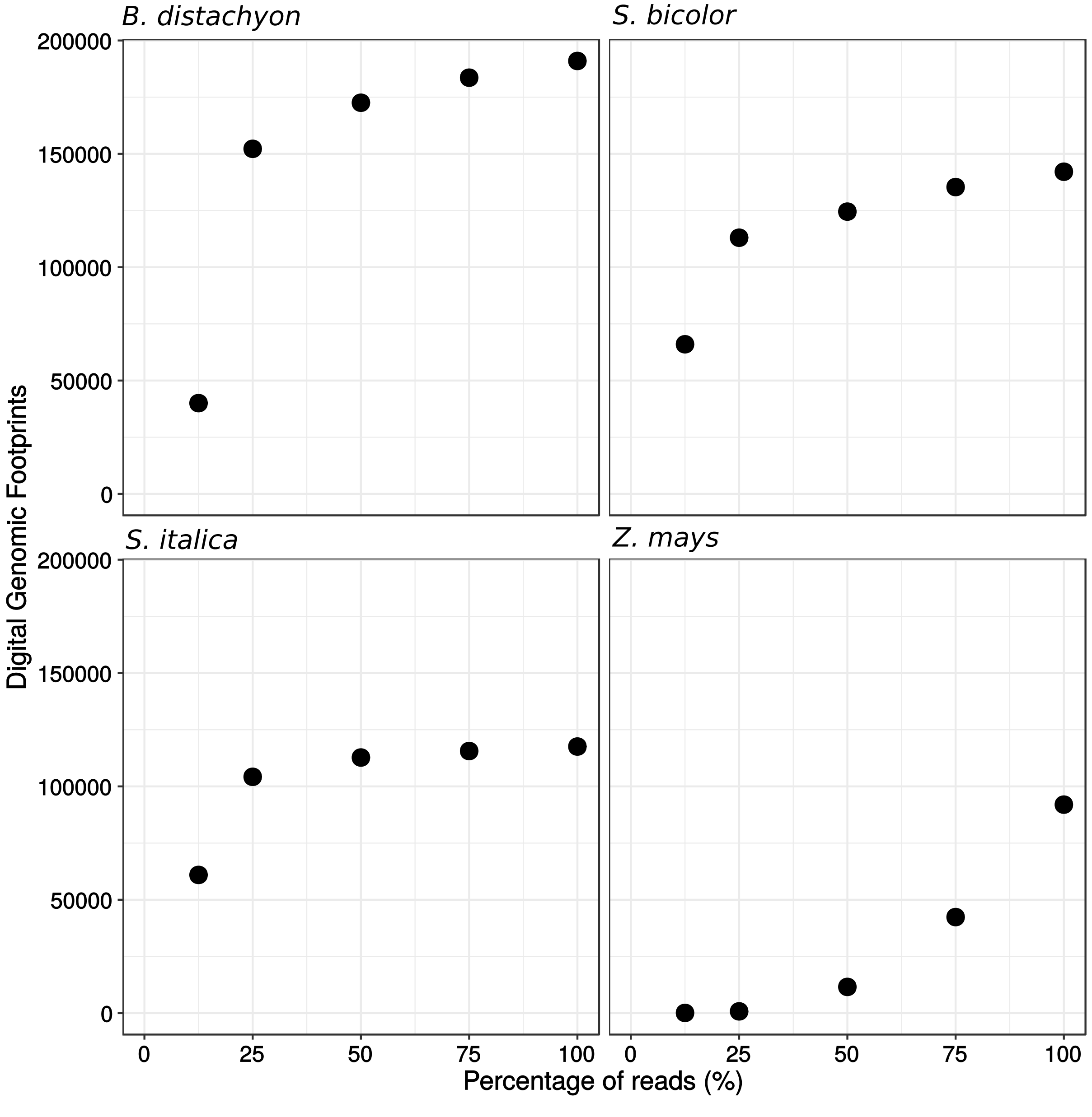


Supplemental Figure 3

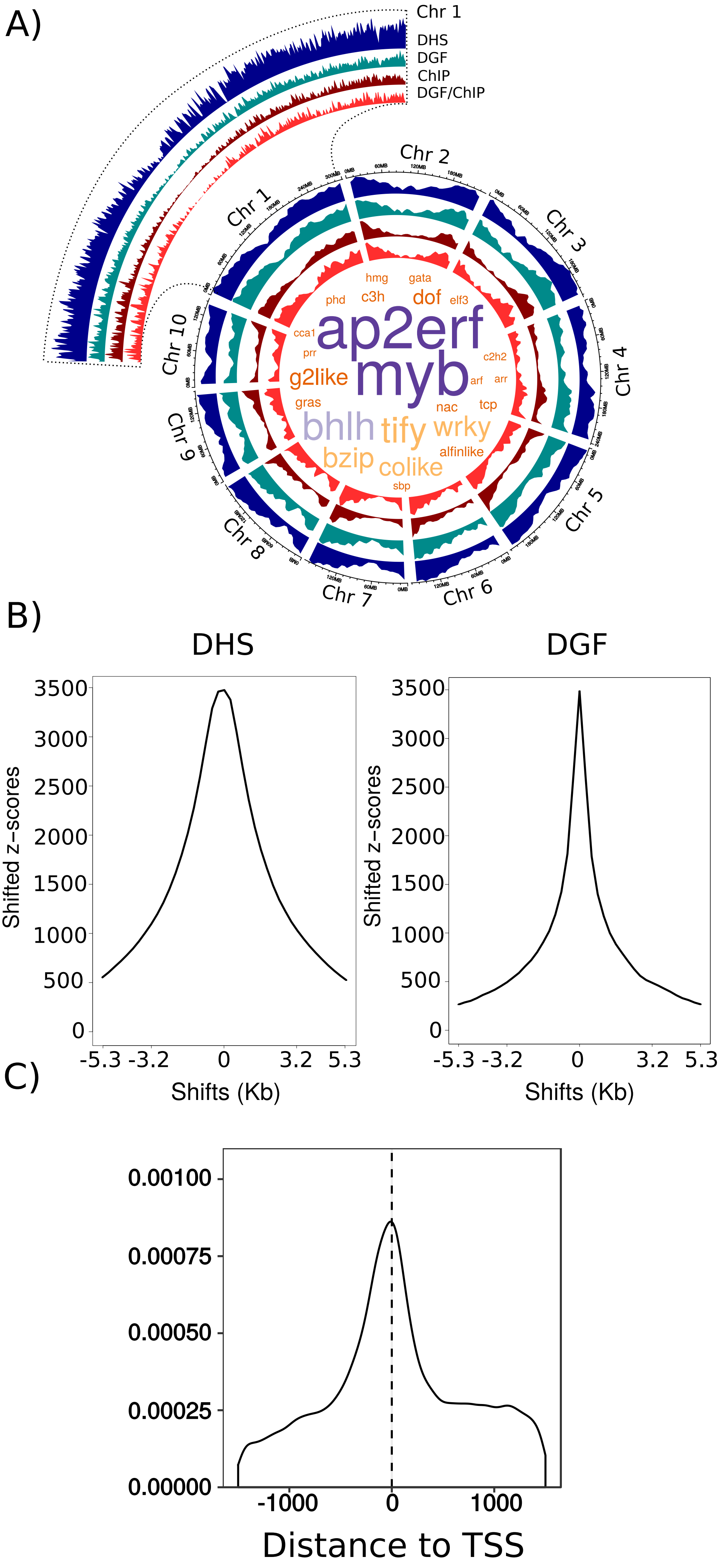


Supplemental Figure 4

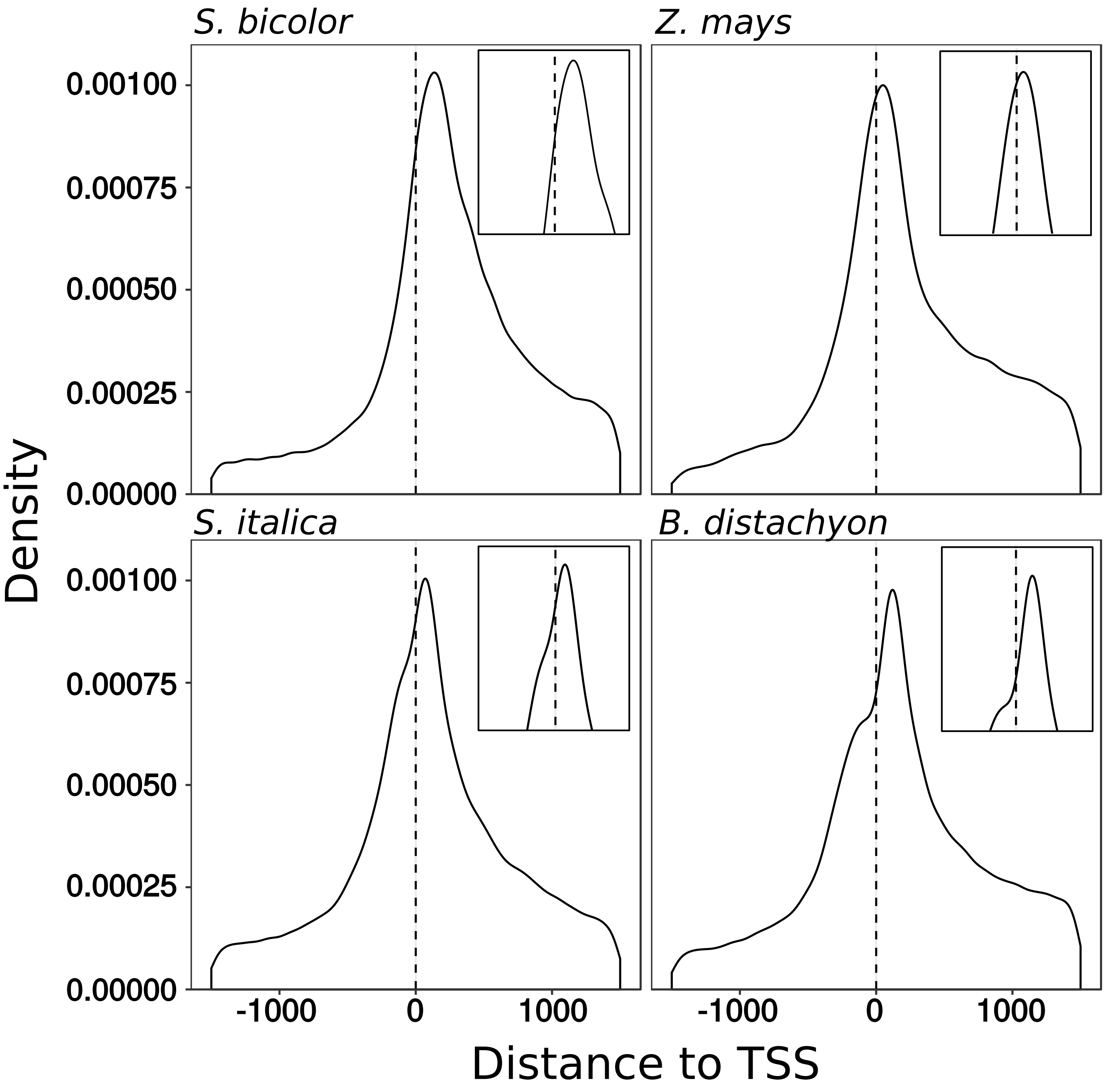


Supplemental Figure 5

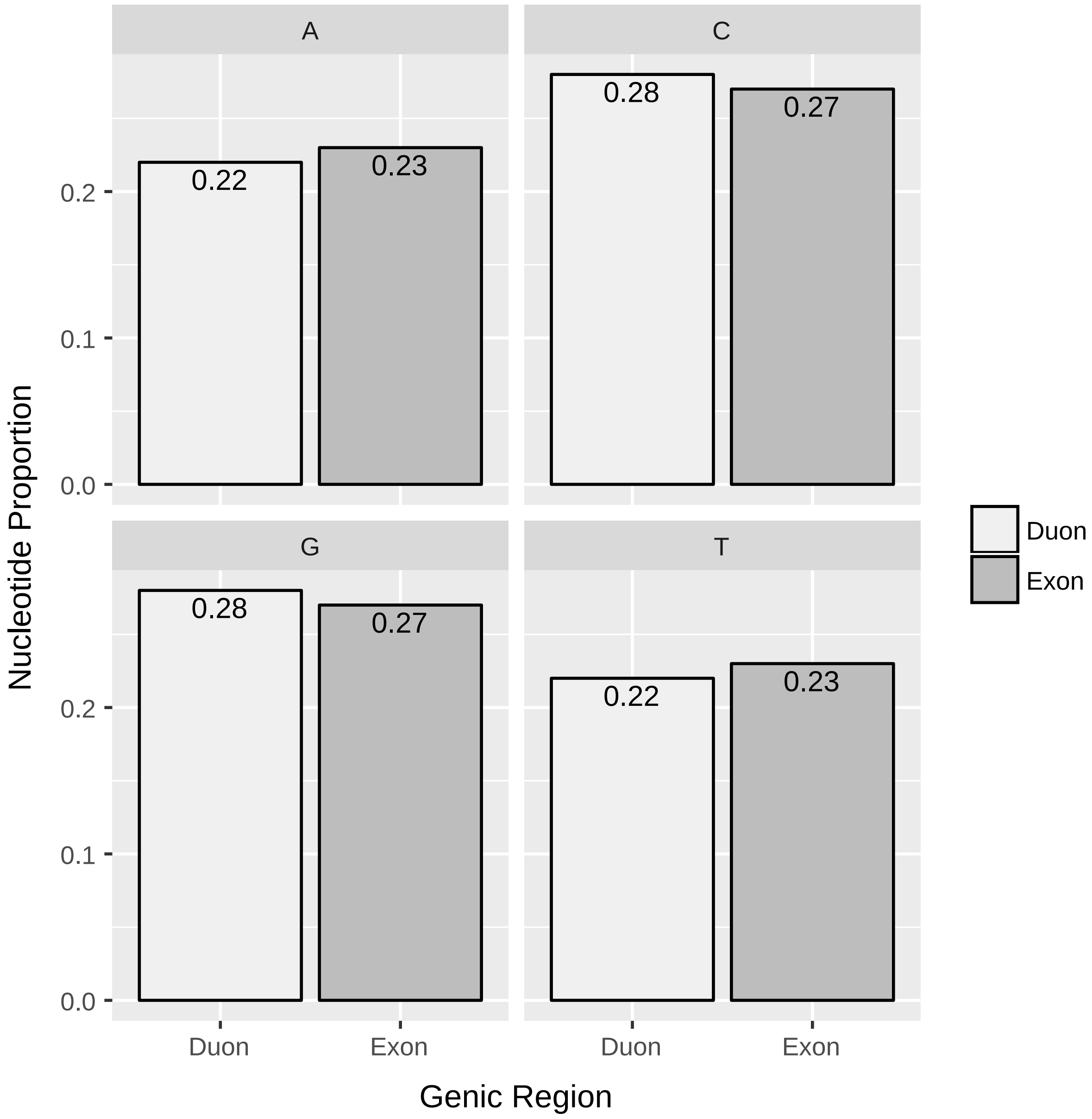


Supplemental Figure 6

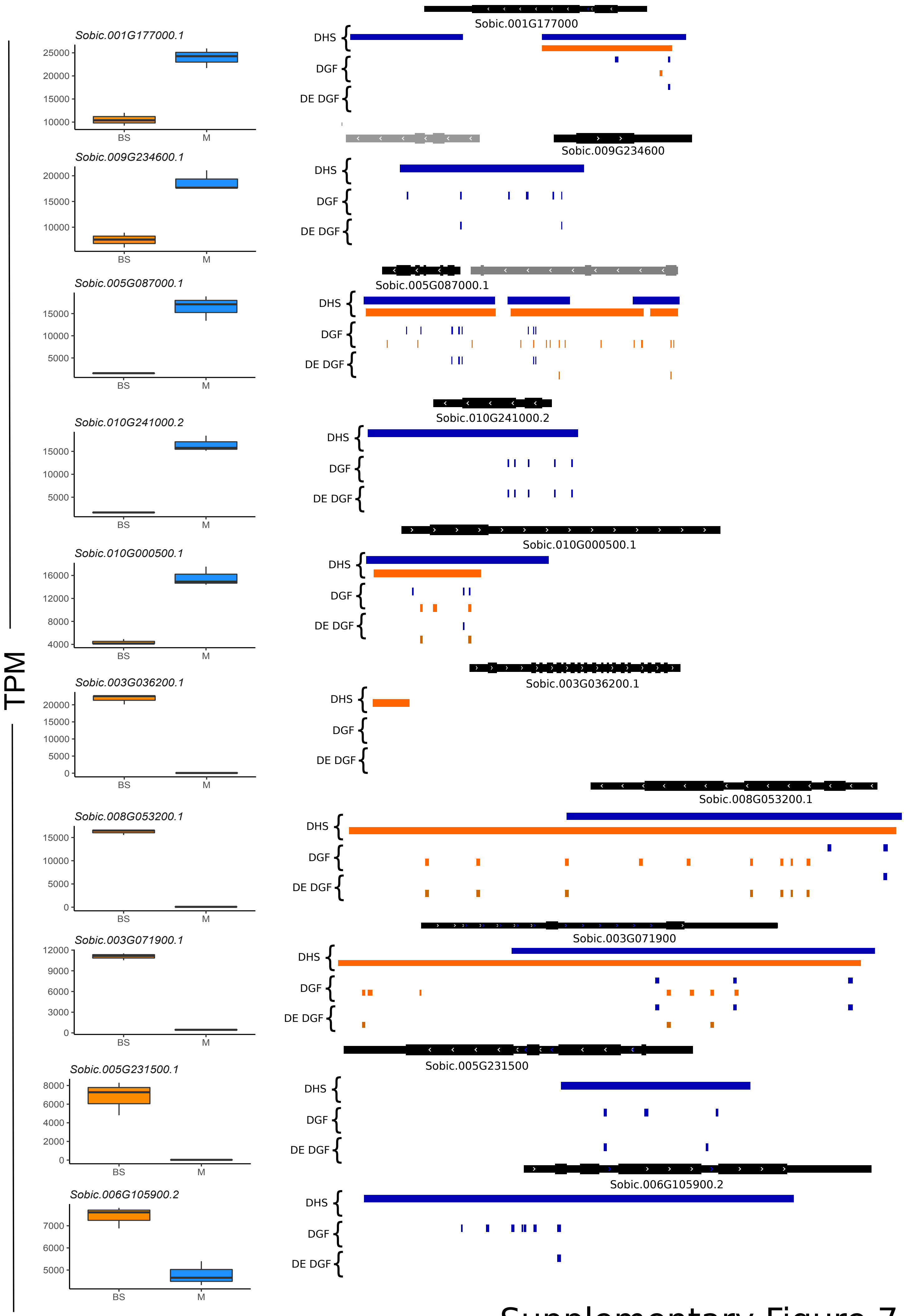


Supplementary Figure 7

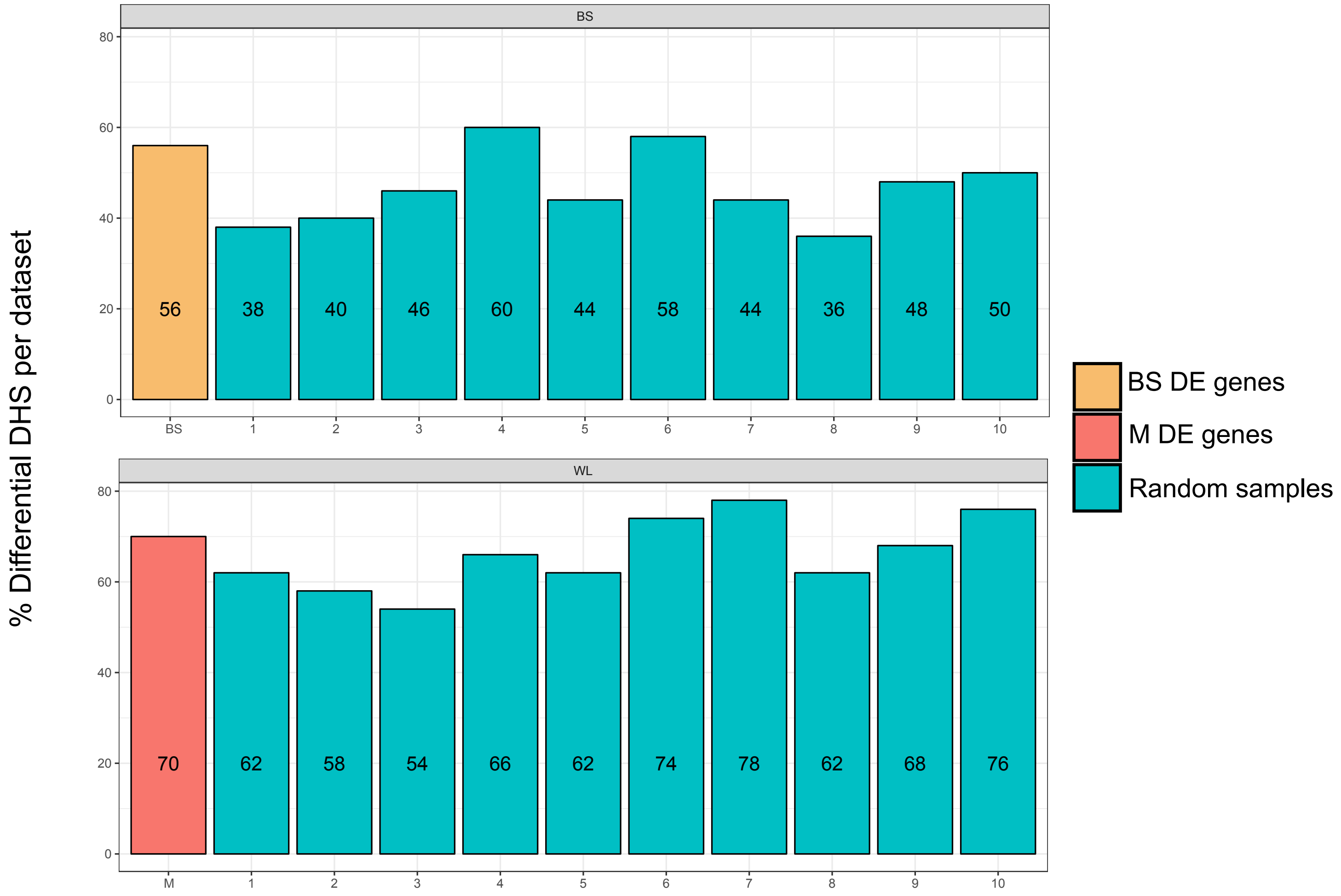


Supplementary Figure 8

#### *S. bicolor*

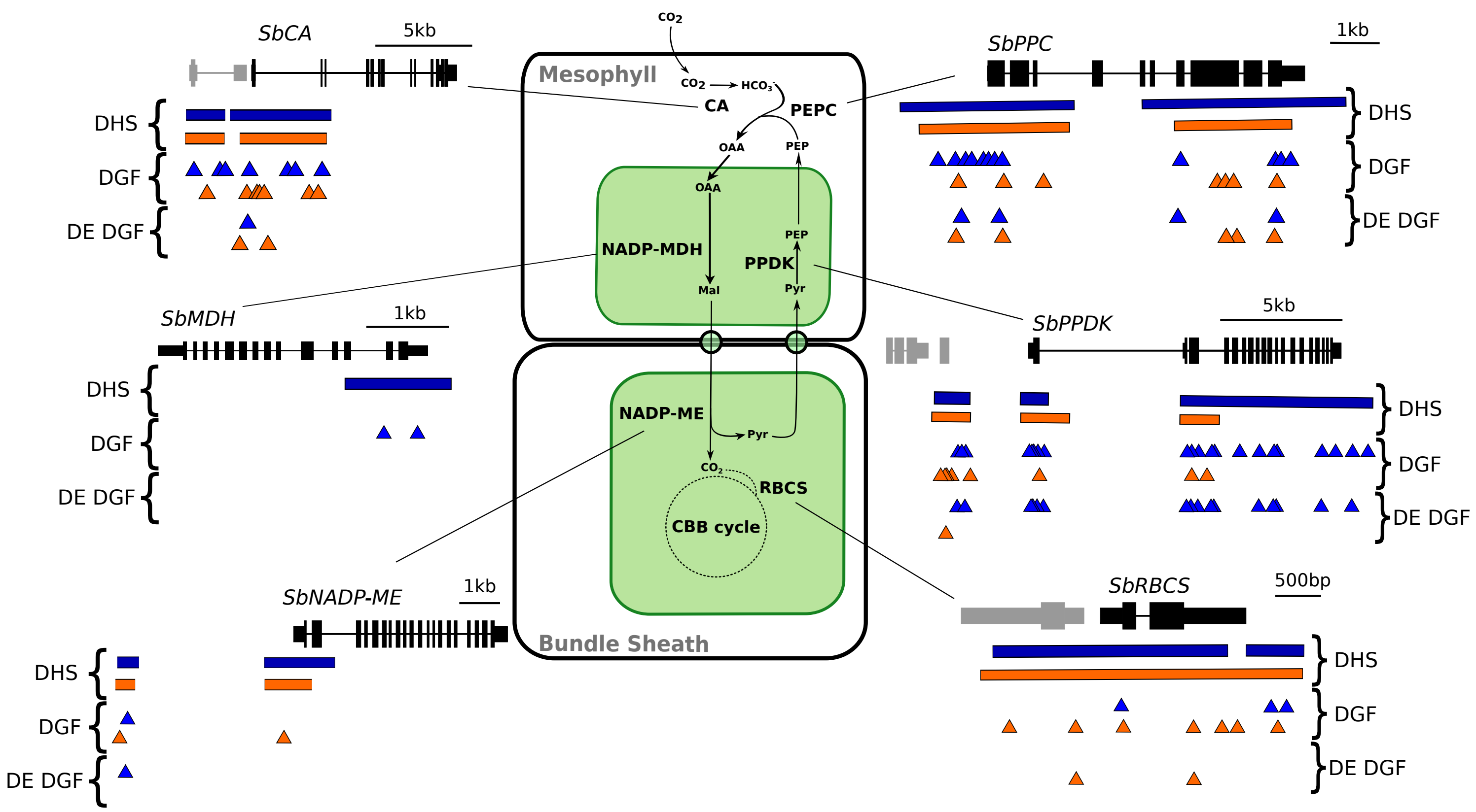


Supplemental Figure 9

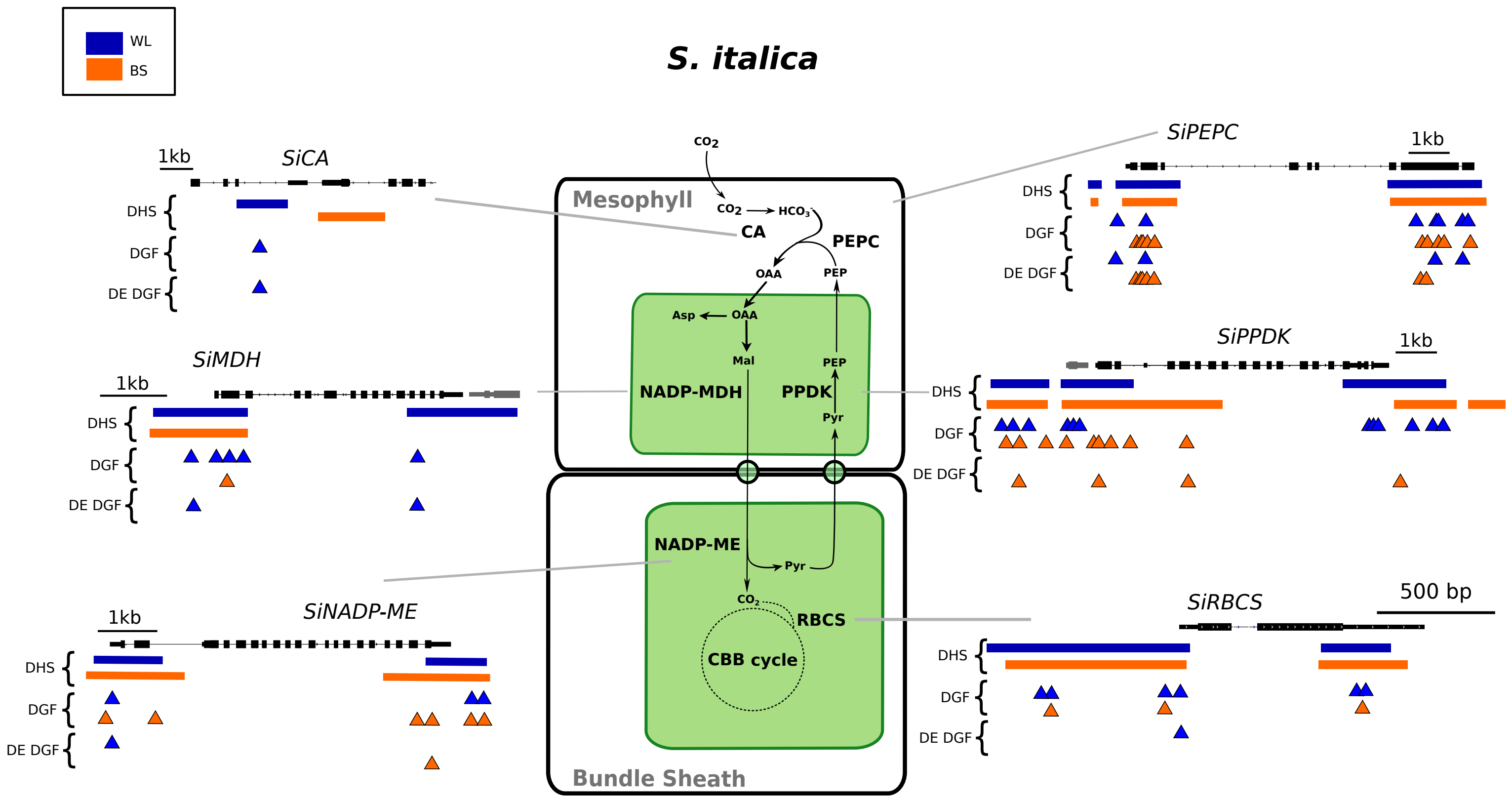


Supplementary Figure 10

### Z. mays

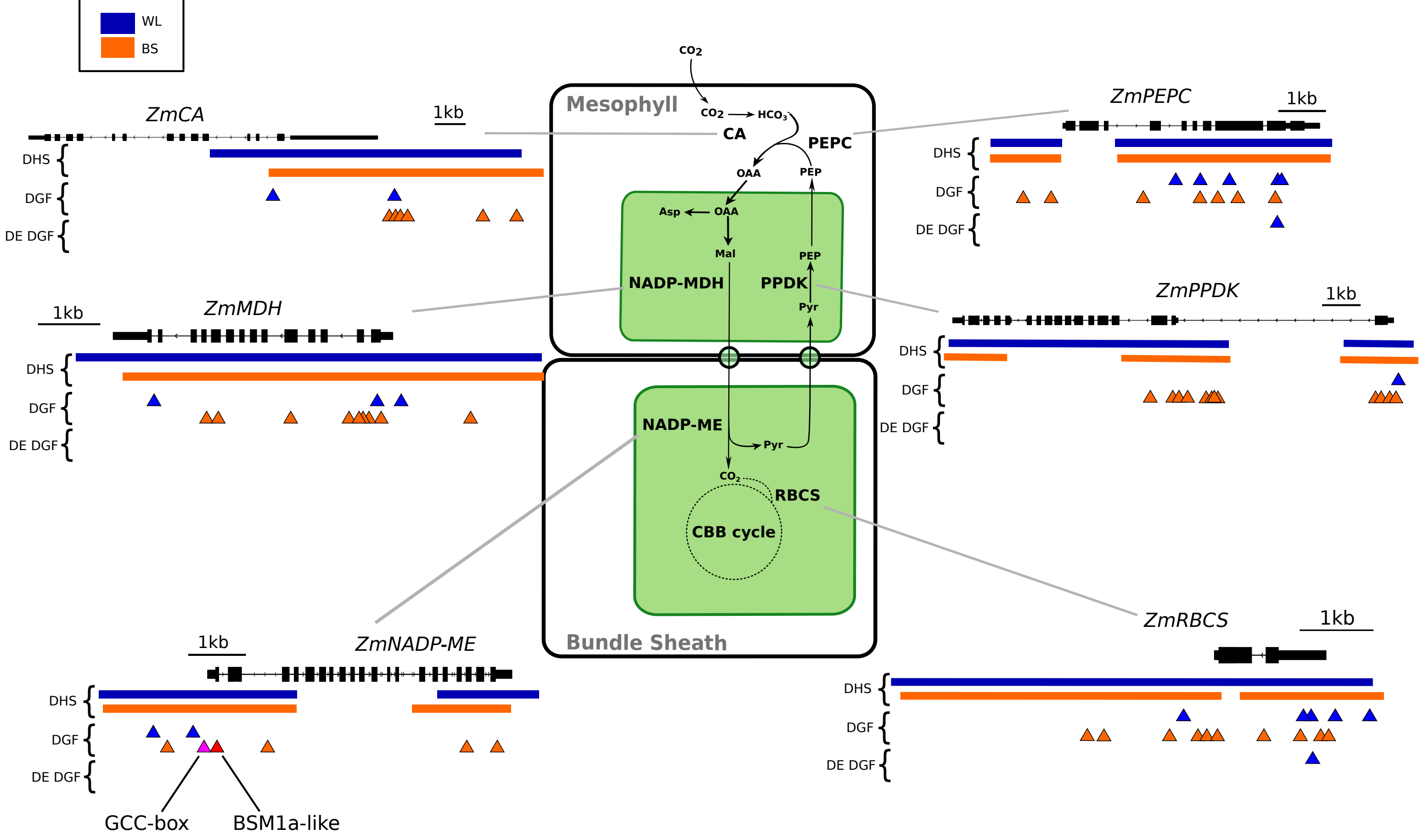


Supplemental Figure 11
